# *Streptomyces* endophytes promote the growth of *Arabidopsis thaliana*

**DOI:** 10.1101/532309

**Authors:** Sarah F. Worsley, Jake Newitt, Johannes Rassbach, Sibyl F. D. Batey, Neil A. Holmes, J. Colin Murrell, Barrie Wilkinson, Matthew I. Hutchings

## Abstract

*Streptomyces* bacteria are ubiquitous in soils and are well-known for producing secondary metabolites, including antimicrobials. Increasingly, they are being isolated from plant roots and several studies have shown they are specifically recruited to the rhizosphere and the endosphere of the model plant *Arabidopsis thaliana*. Here we test the hypothesis that *Streptomyces* bacteria have a beneficial effect on *A. thaliana* growth and could potentially be used as plant probiotics. To do this, we selectively isolated streptomycetes from surface washed *A. thaliana* roots and generated high quality genome sequences for five strains which we named L2, M2, M3, N1 and N2. Re-infection of *A. thaliana* plants with L2, M2 and M3 significantly increased plant biomass individually and in combination whereas N1 and N2 had a negative effect on plant growth, likely due to their production of polyene natural products which can bind to phytosterols and reduce plant growth. N2 exhibits broad spectrum antimicrobial activity and makes filipin-like polyenes, including 14-hydroxyisochainin which inhibits the Take-all fungus, *Gaeumannomyces graminis* var. *tritici*. N2 antifungal activity as a whole was upregulated ~2-fold in response to indole-3-acetic acid (IAA) suggesting a possible role during competition in the rhizosphere. Furthermore, coating wheat seeds with N2 spores protected wheat seedlings against Take-all disease. We conclude that at least some soil dwelling streptomycetes confer growth promoting benefits on *A. thaliana* while others might be exploited to protect crops against disease.

**Importance:** It is vital that we reduce our reliance on agrochemicals and there is increasing interest in using bacterial strains to promote plant growth and protect against disease. Our study follows up reports that *Arabidopsis thaliana* specifically recruits *Streptomyces* bacteria to its roots. In particular, we test the hypothesis that these bacteria can offer benefits to their *A. thaliana* hosts and that strains isolated from these plants might be used as probiotics. We isolated *Streptomyces* strains from surface washed *A. thaliana* roots and genome sequenced five phylogenetically distinct strains. Genome mining and bioassays indicated that all five strains have plant growth promoting properties, including production of IAA, siderophores and ACC deaminase activity. Three strains significantly increased *A. thaliana* growth *in vitro* and when applied in combination in soil. Another produces potent filipin-like antifungal metabolites and we used it as a seed coating to protect germinating wheat seeds against the fungal pathogen *Gaeumannomyces graminis* var. *tritici* (wheat Take-all fungus). We conclude that introducing an optimal combination of *Streptomyces* strains into the root microbiome can provide significant benefits to plants.

## Introduction

The bacterial genus *Streptomyces* comprises more than 600 known species and they produce a diverse array of specialised metabolites that account for ~55% of the antibiotics currently used in human medicine (1). They are filamentous, spore-forming bacteria that are ubiquitous in soils where they play an important role in breaking down complex organic material (2, 3). Intriguingly, they only produce around 10% of their encoded secondary metabolites *in vitro* (2–4). Thus, understanding the role and regulation of their specialised metabolites in natural habitats is essential if we are to unlock the other 90% and discover new molecules (3). Increasingly, *Streptomyces* species are being recognised as important defensive symbionts of a wide range of invertebrate species including bees, beetles, digger wasps and ants (5–9). In addition to this, streptomycetes have also been shown to interact extensively with plant roots, inhabiting both the soil surrounding the plant root, called the rhizosphere, as well as the niche within and between root cells, referred to as the endophytic compartment (10, 11). It has even been suggested that their filamentous, hyphal growth and complex specialised metabolism may have evolved to facilitate interactions with plant roots, presumably to allow entry into root tissue and enable them to compete for food in the form of root exudates or more complex polymers that make up the plant cell wall (2).

Several recent studies have reported that streptomycetes are present, and sometimes enriched, in the endophytic compartment of the model plant *Arabidopsis thaliana* relative to the bulk soil (12–15) where they are attracted by plant metabolites in the root exudates such as salicylate and jasmonate (16, 17). They have also been isolated from the endosphere of many other plant species, including wheat, a crop of huge social and economic value (18–21). Due to their capacity to produce a large number of antimicrobial compounds and their ability to abundantly colonise plant roots, streptomycetes are gaining increasing interest from a biocontrol point of view (10, 11, 19, 22). A recent study demonstrated that certain strains can act as defensive mutualists of strawberry plants whereby they protect their host plant and pollinating bees against fungal infections (23). Many other isolates have been shown to protect important crops against infection and two strains have been developed into commercial biocontrol agents called Actinovate® and Mycostop® (10, 19, 22, 24). The ubiquity of streptomycetes in soil and their diverse specialised metabolism, combined with their ability to colonise plant roots, makes streptomycetes attractive for this purpose. Their spore-forming capabilities also makes them tolerant of many environmental pressures, allowing them to be applied as dried seed coatings which remain viable under various agricultural conditions.

The aim of this study was to test the hypothesis that plant-associated streptomycetes provide benefits to their host *A. thaliana* plant and that strains isolated from *A. thaliana* might confer benefits to important crop plants, such as wheat. We hypothesised that these strains may play defensive or plant growth promoting roles since this genus is consistently recruited to the plant root microbiome. To this end, we generated high quality genome sequences for five *Streptomyces* species which we isolated from the root microbiome of *A. thaliana* plants and which could recolonise *A. thaliana* roots when applied as seed coatings. All five strains, named L2, M2, M3, N1 and N2, encode a large number of secondary metabolite biosynthetic gene clusters (BGCs) and all strains inhibited at least one bacterial or fungal pathogen. Strain N2 demonstrated broad spectrum antifungal and antibacterial activity and was able to inhibit growth of the Take all fungus, an economically important pathogen of wheat, both *in vitro* and on germinating wheat seeds. The antifungal activity of N2 was increased 2-fold *in vitro* in response to indole-3-acetic acid (IAA) and purification of the antifungal molecules identified a number of filipin-like compounds. Curiously, N2 reduced *A. thaliana* growth *in vitro* (but not in soil) which may have been caused by filipins targeting sterols in the plant cell membranes at high concentrations. Strains L2, M2 and M3 all promoted *A. thaliana* growth *in vitro* and when applied in combination to seeds planted in soil, and they all have well characterised plant growth promoting (PGP) traits including the production of plant growth hormones, siderophores and ACC deaminase. We conclude that *A. thaliana* can acquire significant benefits from recruiting streptomycetes to their root microbiome which they likely attract through the deposition of root exudates and dead root material into the bulk soil. Additionally, mining plant-streptomycete interactions may yield novel biocontrol and plant-growth promoting agents that could be developed for future applications in agriculture.

## Results

### Isolation and genome analysis of *A. thaliana* root-associated *Streptomyces* strains

To culture bacteria from *A. thaliana* roots, the roots were washed and sonicated in sterile buffer (as described in 12) to remove soil particles. This process removed all bacteria apart from those there were tightly bound to the plant root surface (the rhizoplane) or were living within the plant root tissue as endophytes (12). The roots were then crushed in sterile 10% glycerol and serial dilutions were plated onto soya flour mannitol, starch casein and minimal salts agar. Colonies resembling streptomycetes were purified by restreaking and were then identified by colony PCR and 16S rRNA gene amplicon sequencing using the universal primers PRM341F and MPRK806R (Table S1). Based on 16S rRNA sequencing, five phylogenetically distinct strains (L2, M2, M3, N1 and N2) were then selected for genome sequencing. We used the PacBio RSII platform (as described in 25) to generate high-quality genome sequences for the five isolates, in addition to three strains of *Streptomyces lydicus*, which are known plant endophytes (26, 27). One of these strains was isolated from the horticultural product Actinovate® and the other two came from the ATCC culture collection (Table S1). All eight linear genomes are within the size range typical for this genus and do not show any significant reductions compared to the genomes of other sequenced *Streptomyces* species (Table 1). The genomes of the *A. thaliana* associated strains L2, M2, M3, N1 and N2 were uploaded to the automated multi-locus species tree (28) for phylogenetic classification. The highest average nucleotide identity (ANI) values of strains L2, M2 and M3 to strains in the database are 88.3%, 94.7% and 91.1%, respectively; these are below the 95% threshold that is generally used to assign strains to a known species (29, 30) so they could be new *Streptomyces* species. Strain N1 has a 98.7% ANI to *Streptomyces albidoflavus* and strain N2 has a 97.6% ANI to *Streptomyces griseofuscus*, suggesting they belong to these clades (Table 1).

**Table 1.**
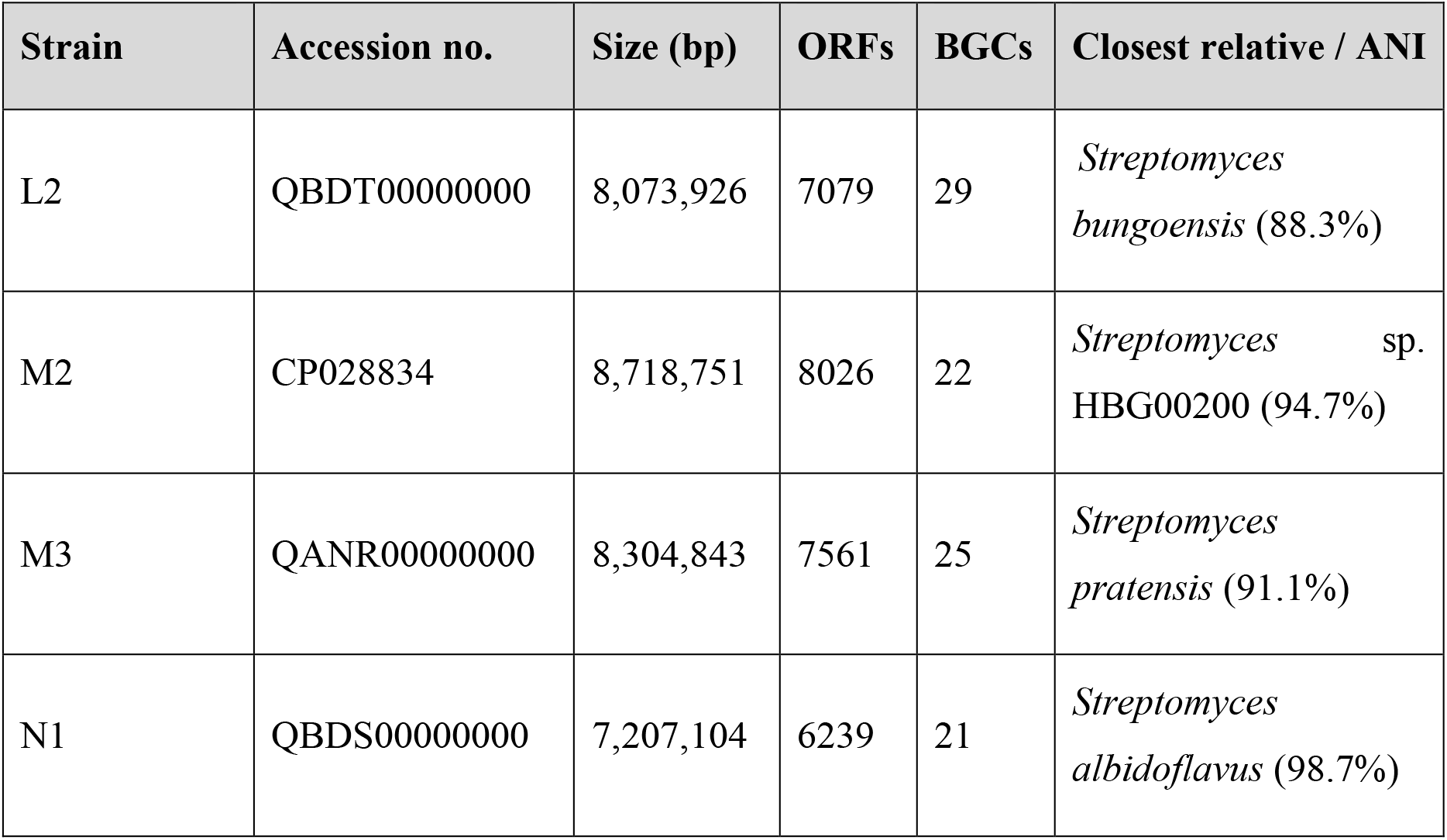
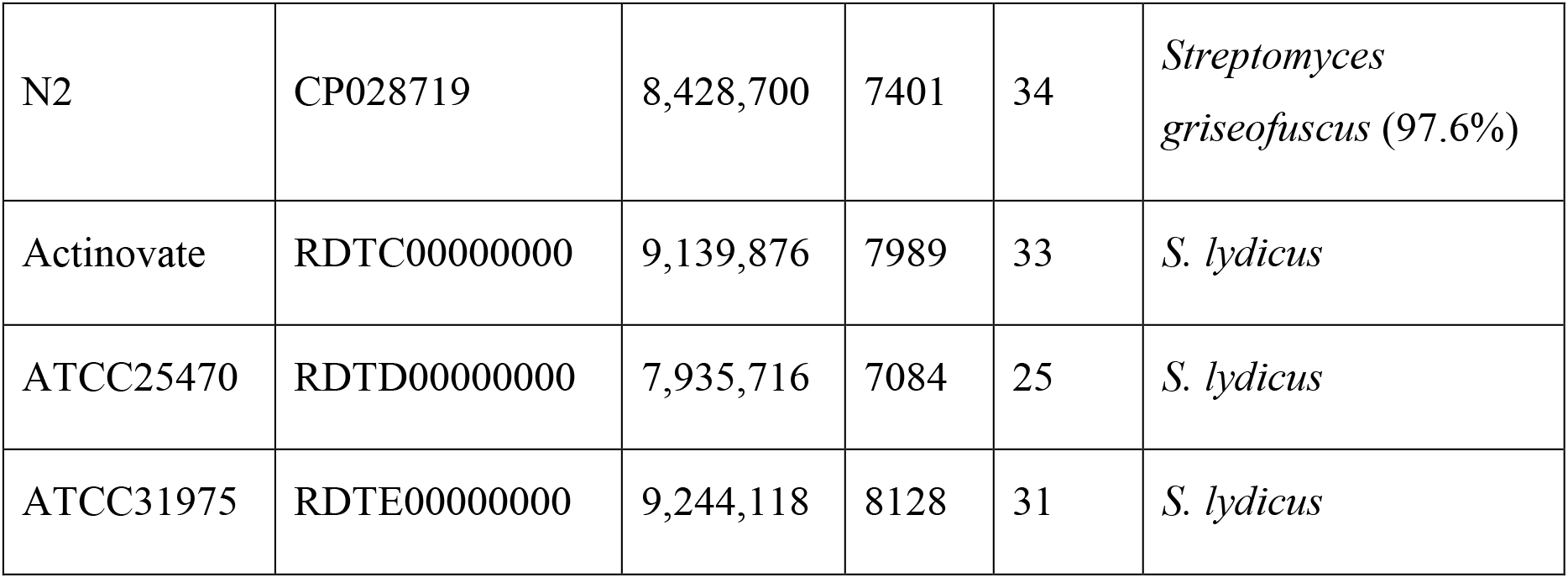
Genome features of root-associated *Streptomyces* strains sequenced for this study, including their GenBank Accession numbers, their genome size in base pairs (bp) and the total number of open reading frames (ORFs) per genome. Genomes were sequenced using the PacBio RSII platform. Biosynthetic Gene Clusters (BGCs) were predicted using AntiSMASH 5.0 (31) and % ANI and closest relatives were determined using AutoMLST (28).

### *Streptomyces* bacteria colonize *A. thaliana* roots

To investigate whether the sequenced *Streptomyces* strains can promote plant growth and fitness, we established root infection assays in which seeds were coated with a suspension of pre-germinated *Streptomyces* spores. Tagging the strains with eGFP and the apramycin resistance (*aac*) gene allowed visual confirmation of root infection using confocal microscopy (Fig. 1) and selective re-isolation of the strains on agar plates containing apramycin, confirming that they were able to re-colonise plant roots.

**Figure 1.**
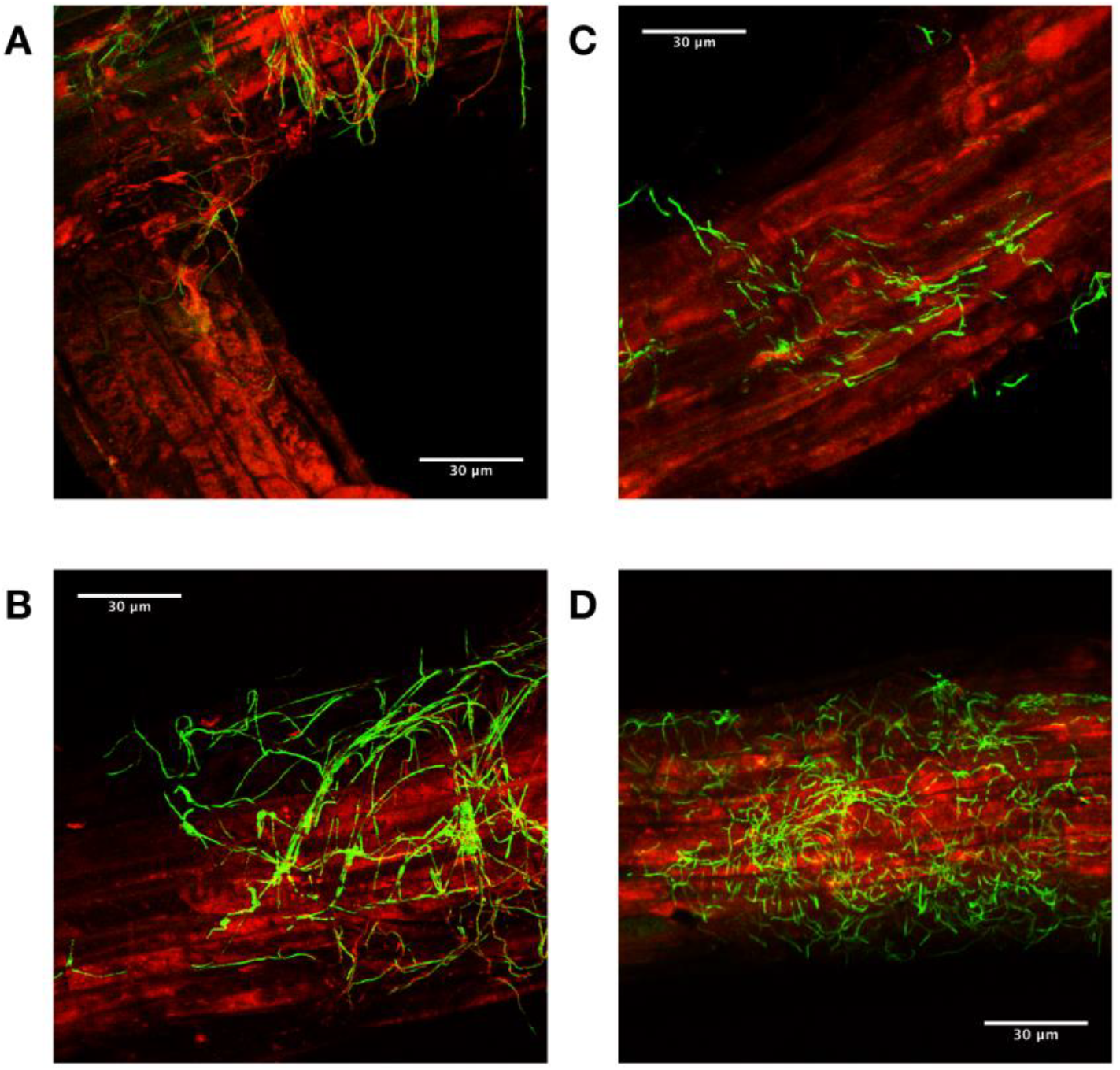
Confocal Laser Scanning Microscopy images of *A. thaliana* rhizoplane colonisation by eGFP-tagged *Streptomyces* strains three days after inoculation. A and B show *A. thaliana* roots (red) colonised by eGFP-tagged *Streptomyces coelicolor* M145 (green) which is a known root endophyte and was used as a control (32). C and D show *A. thaliana* roots (red) colonised by eGFP-tagged *Streptomyces* strain M3 which was isolated in this study (green).

### *Streptomyces* strains M2, M3 and L2 have growth-promoting effects in *A. thaliana*

Next we wanted to determine if any of the *Streptomyces* strains isolated from *A. thaliana* roots can enhance plant growth, so we established root infection assays on agar plates, whereby *Streptomyces* spores were inoculated directly onto the roots of young *A. thaliana* seedlings. We tested all the genome-sequenced strains from this study and found that the inoculation of different strains had a significant effect on the dry weights of plants grown on agar, compared to a sterile control (Fig. 2, F_(8,135)_= 27.63, *P* < 0.001 in an ANOVA test). Strains L2, M2 and M3 significantly increased *A. thaliana* dry biomass under these conditions when compared to sterile control plants (Fig. 2, *P* < 0.05 in Tukey’s Honestly Significant Difference (HSD) tests). In comparison, the three strains of *S. lydicus* had no effect on *A. thaliana* plant biomass *in vitro* despite this species being previously noted to have plant-growth-promoting effects (26). *Streptomyces* strains N1 and N2 significantly reduced the growth of *A. thaliana in vitro* relative to control plants (Fig. 2; *P* < 0.05 in Tukey’s HSD).

**Figure 2.**
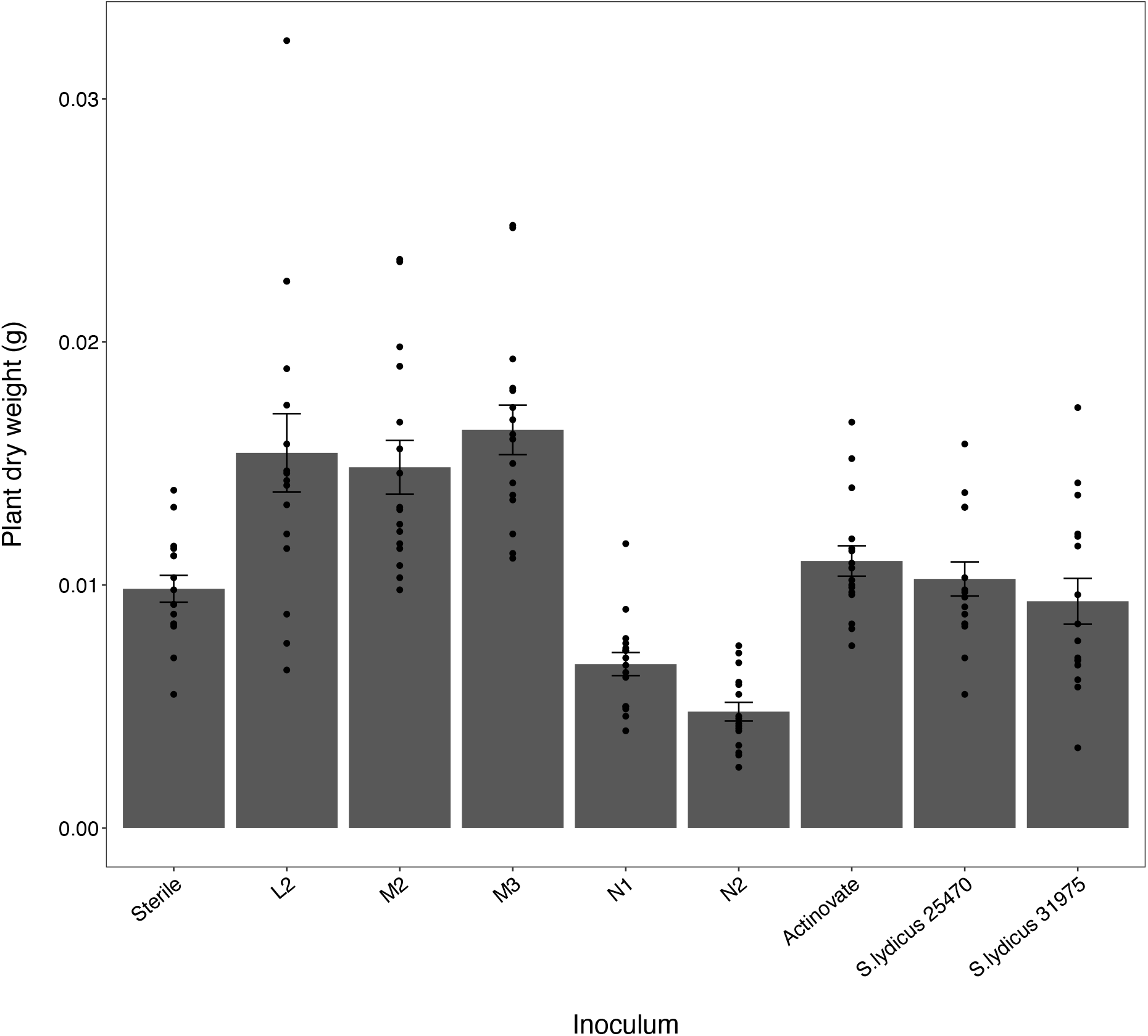
The average biomass of *Arabidopsis thaliana* plants growing on agar plates, following inoculation with sequenced *Streptomyces* isolates. Biomass (dry weight in grams) was measured 16 days after inoculation. Sterile plants were grown as a control. N=16 plants per treatment, represented by points. Error bars represent standard errors.

To test whether each of the new isolates could promote plant growth in soil, spores of each strain were applied individually to *A. thaliana* seeds planted in Levington’s F2 seed and modular compost. Since, it is known that some strains can work synergistically to promote plant growth (33–35), a mixture of spores of L2, M2 and M3 (the strains that promoted plant growth *in vitro*) was also added to seeds. Strain inoculation had a significant influence on the dry weight of plants grown in compost (ANOVA test on log-transformed dry weight: F_(7,58)_ = 3.9358, *P* = 0.001). However, interestingly none of the strains had an effect on plant growth when applied individually (Fig. 3, *P* >0.05 in Tukey HSD test), but the application of a spore mixture of L2, M2 and M3 significantly increased plant dry weight from an average of 12.69 mg ± 1.94 (mean ± SE) for control plants, to 39.29 mg ± 4.39 (Fig. 3).

**Figure 3.**
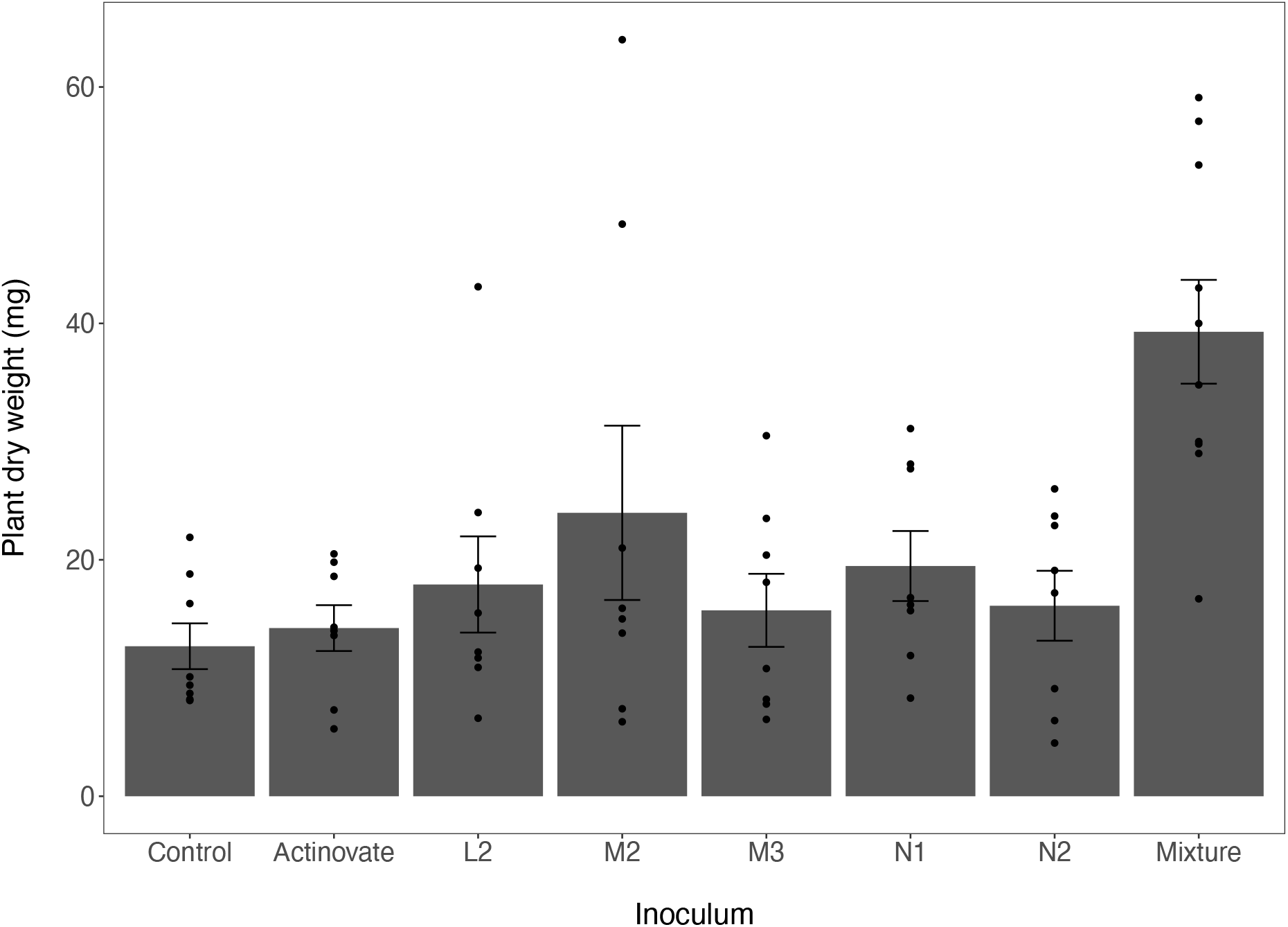
Total dry weight of *A. thaliana* plants grown in Levington’s F2 compost from seeds inoculated with spores of Actinovate, L2, M2, M3, N1 and N2 or a mixture of L2, M2 and M3 *Streptomyces* spores. Dry weight is shown in milligrams. Sterile seeds were grown as a control. N=8 replicate plants per treatment (shown as points).

### Characterisation of plant growth promoting traits

KEGG pathway analysis revealed that the genomes of all our sequenced *Streptomyces* strains possess genes encoding proteins involved in the biosynthesis of IAA, which can contribute to shoot and root growth (Table S2). For example, all strains have genes encoding key proteins involved in the indole-3-acetamide (IAM) pathway, whereby tryptophan is converted to IAM via a tryptophan 2-monooxygenase enzyme (KEGG reaction R00679). IAM is then further converted to IAA through the action of an amidase enzyme (KEGG reaction R03096). Several strains also possessed genes encoding enzymes involved in the tryptamine (TAM) pathway. *In vitro* colorimetric assays using Salkowski reagent (as described in 36) qualitatively confirmed the ability of all strains to make IAA (Fig. S1). In addition to IAA, the genomes of all the *Streptomyces* isolates possess up to two copies of genes encoding the enzyme aminocyclopropane-1-carboxylate (ACC) deaminase. This cleaves ACC, which is the direct precursor for the plant phytohormone ethylene, into ammonia and 2-oxobutanoate, Kegg reaction: R00997 (Table S2). Bacteria can use the ammonium released in this reaction as a nitrogen source, and all the isolates are capable of utilising ACC as a sole nitrogen source in minimal medium (Fig. S2). There is evidence that the activity of the ACC deaminase enzyme can reduce plant damage and early-onset senescence caused by excessive ethylene production under prolonged periods of plant stress, by removing the substrate for ethylene biosynthesis (37, 38).

### Bioactivity of root-associated *Streptomyces* isolates

*Streptomyces* bacteria are well-known for their ability to produce a wide range of specialised metabolites which can have bioactivity against bacteria, fungi, viruses, nematodes, insects and plants (1, 3, 39). We reasoned that our strains likely make antimicrobial natural products and that strains N1 and N2 may encode herbicidal compounds given that they have an adverse effect on *A. thaliana* growth *in vitro* (Fig. 2). All eight sequenced genomes were submitted to the bacterial antiSMASH 5.0 portal (31) which can predict BGCs for major types of specialised metabolite. This identified between 21 and 34 putative specialised metabolite BGCs in the sequenced genomes, which is within the typical range for this genus (Table S1). This includes BGCs predicted to encode polyketide synthases (PKS), non-ribosomal peptide synthases (NRPS), ribosomally encoded post translationally modified peptides (RiPPs) and terpenes (Table S3). All five strains encode multiple siderophore BGCs which are molecules that chelate metal ions, such as iron, generating soluble complexes that can be taken up by plant roots and contribute to plant growth (40). Siderophores also help microorganisms to compete in the soil, rhizosphere and endosphere, with the added benefit that this may act to exclude plant pathogens that are also competing for iron (41).

To test whether antimicrobial compounds could be produced *in vitro,* sequenced strains were tested for their ability to inhibit a range of bacterial and fungal pathogens, including the bacterial plant pathogen *Pseudomonas syringae* and the fungus *G. graminis* var. *tritici* which is the causative agent of wheat Take-all, one of the most economically damaging diseases of wheat worldwide (19). All eight strains exhibited antibacterial activity and N2 inhibited all of the bacterial strains that were tested: *Bacillus subtilis*, methicillin resistant *Staphylococcus aureus* (MRSA), *Escherichia coli* and *Pseudomonas syringae* (Fig. S3; Table S4). The N2 genome encodes a rich repertoire of BGCs including several putative antibacterial BGCs predicted to encode the proteins responsible for the biosynthesis of albusnodin, albaflavenone and diisonitrile antibiotic SF2768-like antibiotics, the antiproliferative actinomycin D which is used clinically as an anti-cancer therapeutic, in addition to a possible analogue of the proteasome inhibitor cinnabaramide (Table S3). The actinomycins exhibit broad-spectrum bioactivity by binding to DNA and inhibiting transcription; thus it could result in the inhibition of all the pathogenic indicator strains (42). M2 is interesting because it inhibits *Pseudomonas syringae* but none of the other bacteria that were tested (Table S4). It encodes a relatively modest number of BGCs with one for a putative aquamycin-type antibiotic (Table S3). However, this effect may be due to a siderophore because an antibacterial compound that inhibits *Pseudomonas* would also be expected to inhibit *B. subtilis* and *E. coli* (Table S4). L2 encodes clusters for the biosynthesis of albaflavenone and thioviridamide-like molecules (Table S3) and inhibits *B. subtilis in vitro* (Table S4). M3 inhibits *B. subtilis* and *P. syringae* (Table S4) and encodes a type 3 PKS for the biosynthesis of alkylresorcinol-type phenolic lipids (Table S4) which may have antibacterial activity. Finally, N1 inhibits *B. subtilis* (Table S4) and encodes several BGCs encoding molecules with potential antibacterial activity including a putative polycyclic tetramate macrolactam (PTM) antibiotic which are widely distributed but often cryptic (43), and a desotamide like antibiotic (Table S3). It should be noted that up to 90% of specialised metabolite BGCs are cryptic (i.e. not expressed under laboratory conditions) in *Streptomyces* species, largely because we do not understand the environmental or host-derived signals that activate their expression (3). In addition, there are many known specialised metabolites for which the BGCs have not yet been identified so it is likely that at least some of the known BGCs in these genomes are not expressed under the growth conditions used in this study and/or that these strains encode antimicrobials whose BGCs cannot be predicted by antiSMASH 5.0.

The strains L2, M2 and M3 do not exhibit antifungal activity against *C. albicans*, *L. prolificans* or *G. graminis* var*. tritici in vitro* (Table S4), although it is possible that these strains may encode cryptic antifungal compounds that are not expressed under the conditions used in experiments. For example, M3 encodes a clavam-like cluster which has a similar structure to those that encode the antifungal compound clavamycin (Table S3). In comparison, strains N1 and N2 both have type 1 (T1) PKS gene clusters which are predicted to encode the biosynthesis of known polyene antifungal compounds (Table S3). The N1 T1PKS gene cluster is predicted to encode candicidin and the N2 T1PKS gene cluster is predicted to encode filipin-type antifungals (Table S4). Both of these metabolites inhibit fungal growth by binding to sterols in their cell walls and it is known that filipins can also bind to phytosterols in plant cell membranes and reduce the growth of plant roots at high concentrations (44–46). Thus, it is possible that N1 and N2 are making polyenes *in planta* and this is what caused the reduction in growth observed during *in vitro* inoculation experiments (Fig. 2). These strains did not influence plant growth in compost (Fig. 3) suggesting that the effect was diluted in a non-sterile system, or that the polyenes were not expressed under these conditions. Despite the presence of antifungal-like clusters in the other *A. thaliana* isolates, only N2 exhibited antifungal activity *in vitro* (Table S4). In fact, N2 demonstrated potent and broad-spectrum antifungal activity as it inhibited all three test strains; these were the human pathogenic fungal species *Candida albicans* and multi-drug resistant *Lomentospora prolificans* as well as the plant pathogenic Take-all fungus *Gaeumannomyces graminis* var *tritici* (Table S4).

Production of antifungal compounds likely gives N2 an advantage in the rhizosphere and, consistent with this, N2 antifungal activity against *C. albicans* was shown to increase 2-fold *in vitro* in the presence of indole-3-acetic acid (IAA), as judged by the size of the inhibition zone (Fig. 4). IAA is the precursor to the plant phytohormone auxin which regulates processes involved in growth and development (47). It is also made by many microbial species in the rhizosphere, including the bacterial strains that were isolated in this study (Fig. S1 & Table S2), and has been noted previously for its role in both intra- and inter-kingdom signalling (47–49). Thus, N2 may be driven to produce antifungals in close proximity to plant roots when it is in competition with other microbes.

**Figure 4:**
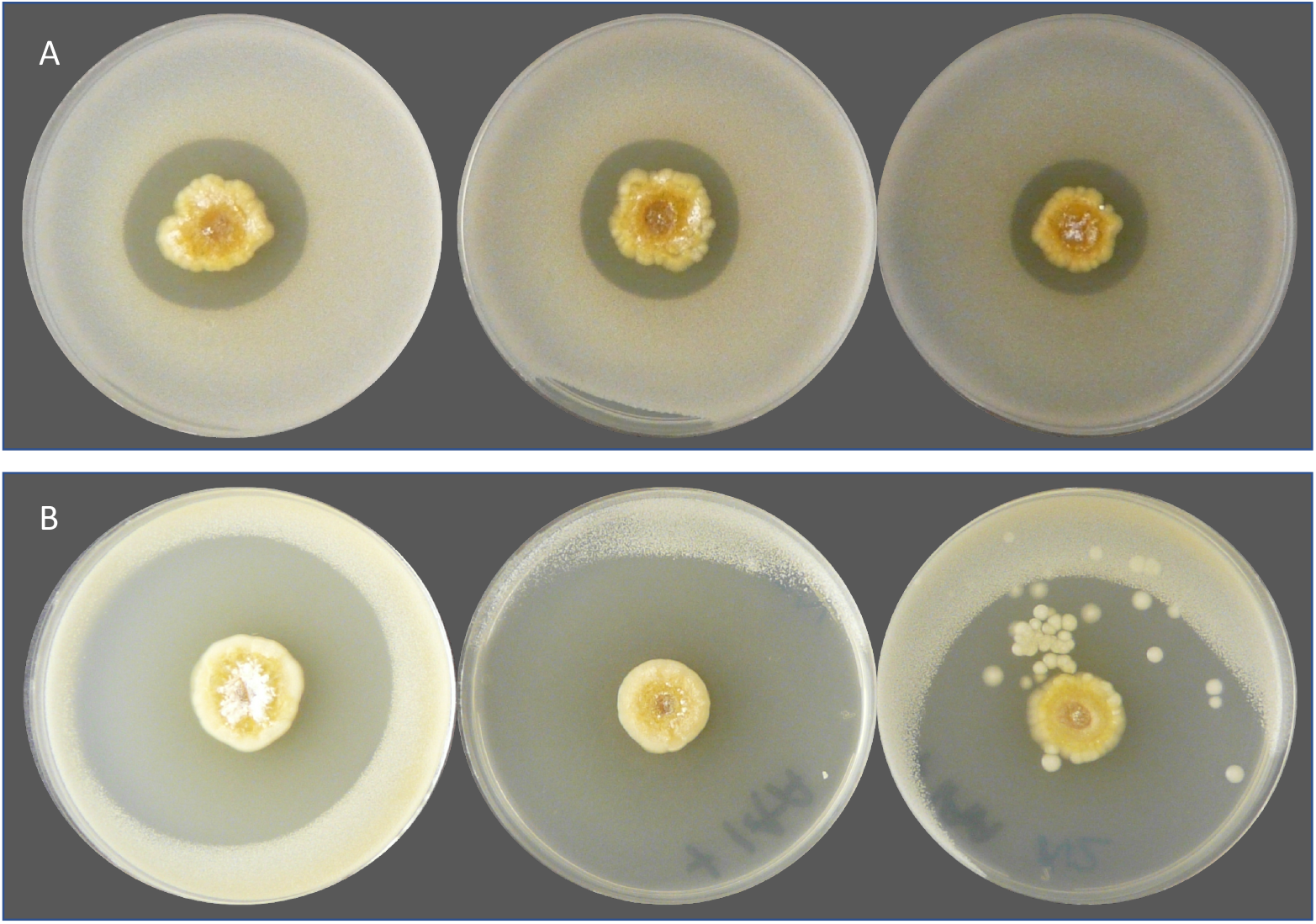
(A) Three biological replicates of strain N2 (centre) growing on minimal medium agar that has been overlaid with soft LB agar inoculated with *Candida albicans.* (B) is the same but the minimal medium agar contains 0.1 mg ml^−1^ of indole-3-acetic acid. This experiment was repeated 4 times (each time with 3 replicates), with consistent results.

### Purification and identification of antifungal compounds from *Streptomyces* strain N2

To extract the molecules with antifungal activity, strain N2 was plated on SFM agar (4 L, 120 plates) and grown for eight days at 30 °C before extraction with ethyl acetate. The crude extract had an intense orange colour and HPLC analysis identified components with UV characteristics typical of polyene metabolites. Comparison to an authentic commercial sample of the filipin complex of *Streptomyces filipinensis* (Sigma Aldrich) confirmed the presence of several filipin related molecules, although the two major components eluted earlier than these molecules (Fig. S4). The two major components eluted very closely together and had *m/z* values consistent with the previously reported molecules pentamycin (also known as fungichromin) and 14-hydroxyisochainin (50–52). Despite a challenging elution profile sufficient separation was accomplished by applying multiple purification steps, and the isolated material enabled structural confirmation by 2D NMR (Figs S5-S16, Tables S5-S7). The extract also contained a mixture of components with an absorption maximum at 444 nm and this mixture was red after separation from the polyene fraction. LCMS analysis indicated that these components were consistent with the known actinomycin congeners D, X_2_ and X_0β_ respectively (Fig. S4). These are most likely encoded by BGC5 in the N2 genome (Table S3) (53). The filipin-like compounds (Fig. 5) can all be assigned to the type 1 PKS BGC in region 1 (Table S3, Fig. S17) (51, 54, 55).

**Figure 5.**
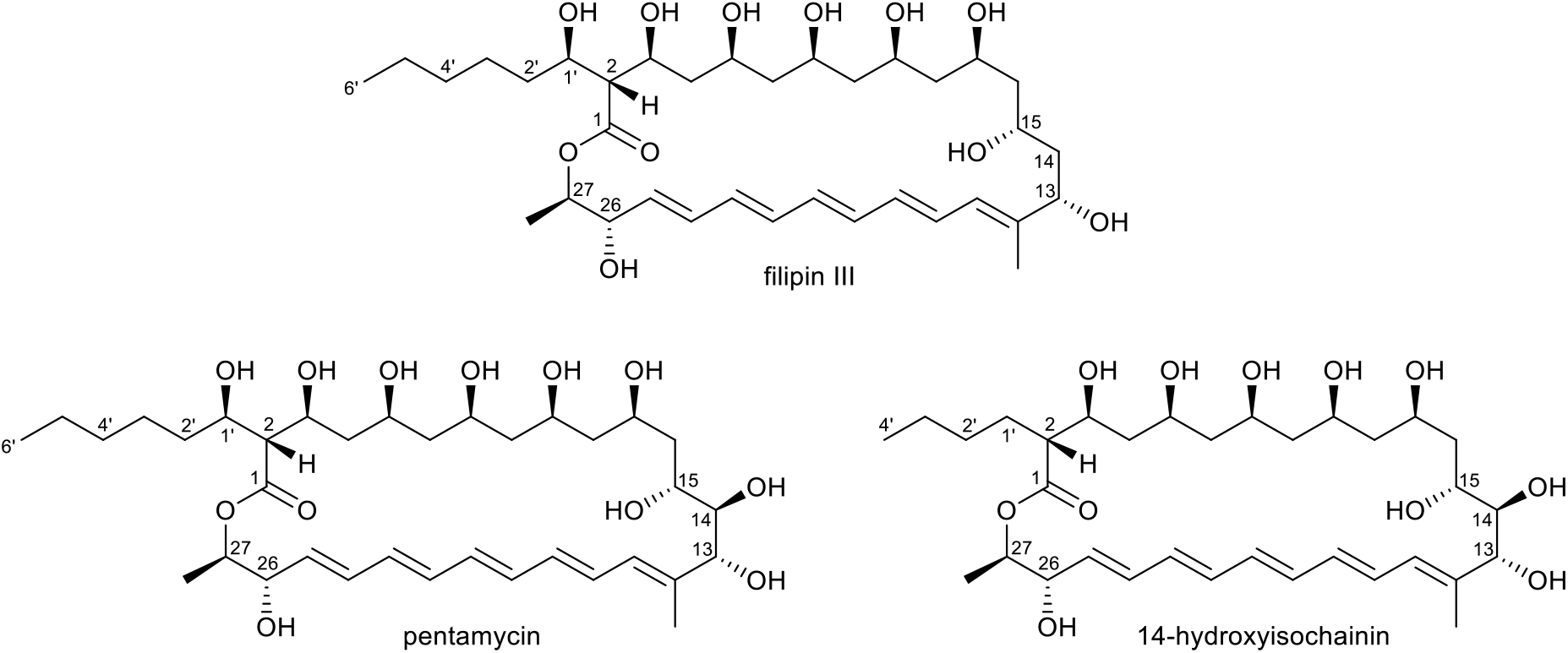
Structures of filipin III, pentamycin and 14-hydroxyisochainin.

Pentamycin is active against *Candida albicans* and *Trichomonas vaginalis* and is used for the treatment of vaginal candidiasis, trichomoniasis, and some mixed infections (56–58). It is identical in structure to filipin III apart from the presence of an additional secondary hydroxyl group at C14 (Fig. 5) which is added by a cytochrome P450 monooxygenase encoded by a gene located directly upstream of the first PKS gene (A1) in the *pentamycin* BGC as reported recently (51); the same BGC architecture is observed in *Streptomyces* strain N2 (Fig. S17). 14-hydroxyisochainin shares the same polyene core structure as pentamycin but carries an altered side chain lacking two carbon atoms and is indicative of different length extender units being utilized by the final module of the type I PKS during biosynthesis (Fig. 5). This observation is unexpected as the co-production of pentamycin and 14-hydroxyisochainin has only been observed as a result of precursor-directed biosynthesis (52). Thus, this example of co-production by *Streptomyces* strain N2 appears to be novel.

Disc diffusion bioassays confirmed that pentamycin and 14-hydroxyisochainin inhibit the growth of *C. albicans* (Fig. S18), but only 14-hydroxyisochainin was able to inhibit *G. graminis* var. *tritici* (Fig. S19), suggesting that 14-hydroxyisochainin is responsible for the antifungal activity of N2 against the Take-all fungus, potentially in combination with lower quantities of other products of the filipin-like BGC. The complex consisting of actinomycin D, X_2_ and X_0β_, which were co-purified from N2 extracts (Fig. S4), did not have antifungal activity suggesting they are not responsible for N2 antifungal bioactivity *in vitro* (Fig. S18). Neither pentamycin, 14-hydroxyisochainin or the actinomycin complex inhibit the growth of *E. coli* K12 (Fig. S18) suggesting that a previously undescribed compound is responsible for the observed inhibition of *E. coli* and *P. syringae* by N2 *in vitro* (Fig. S3, Table S4).

### *Streptomyces* strain N2 protects wheat against Take-all

In order to test whether strain N2 has the potential to protect wheat plants against *G. graminis* var. *tritici,* we inoculated surface sterilised wheat seeds (*Triticum aestivum* var. Paragon) with N2 spores or left them sterile as a control. These seeds were germinated next to a central plug of Take-all fungus on potato glucose agar (PGA) plates. On the control plates, *G. graminis* grew outwards across the agar plate and over the sterile wheat seedlings, whereas the seeds that had been inoculated with N2 spores were resistant to *G. graminis* var. *tritici*, as indicated by a zone of inhibition around the germinating wheat seeds (Fig. 6). The most parsimonius explanation is that the N2 spores have germinated and are producing 14-hydroxyisochainin and possibly other filipins, that inhibit the Take-all fungus.

**Figure 6.**
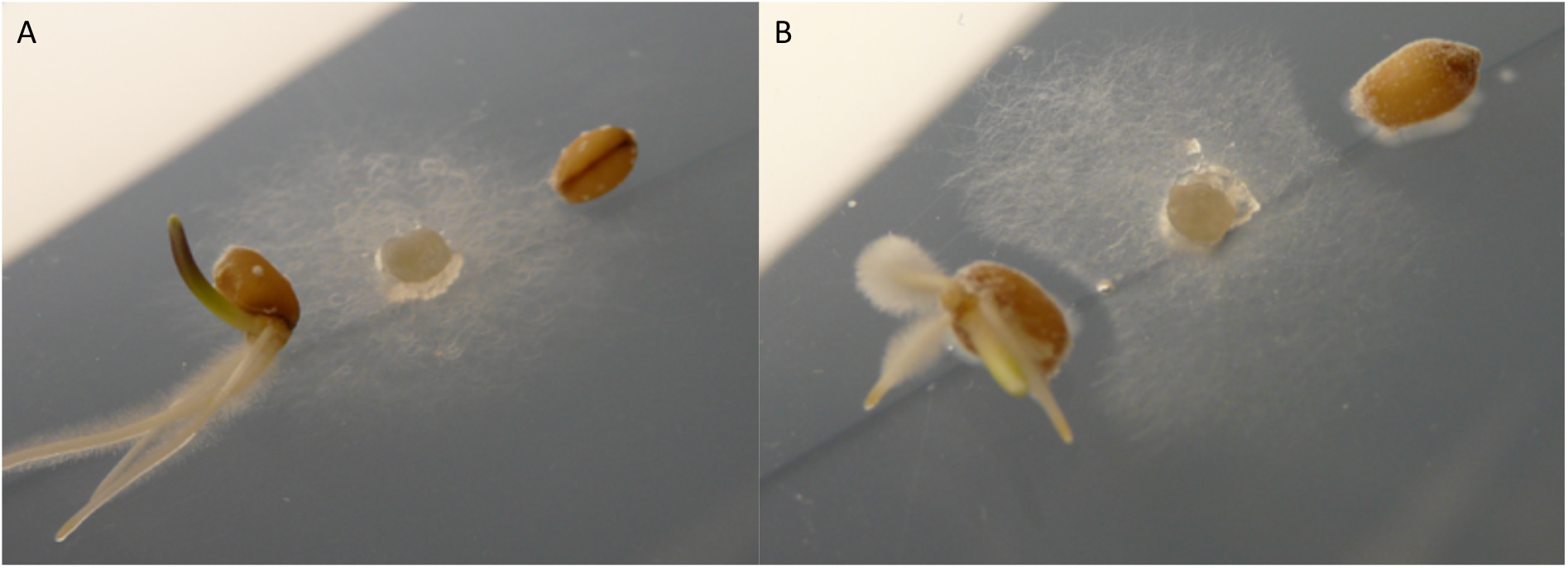
Inhibition of *G. graminis* var. *tritici* in wheat seedlings. Germinating wheat seeds are either A) sterile or B) inoculated with a spore preparation of *Streptomyces* isolate N2, growing next to a plug of *G. graminis* var. *tritici*, the Take-all fungus. *G. graminis* is prevented from growing towards inoculated seeds, as demonstrated by the zone of inhibition.

To further test the potential of the *Streptomyces* strain N2 to act as a biocontrol strain against Take-all *in vivo*, wheat seeds were soaked in N2 spores, allowed to dry, and then grown in sterile vermiculite containing *G. graminis* var. *tritici* mycelia. After 3 weeks of growth at 25°C, Take-all infection severity was scored on a scale of 0-8, using an infection scoring system as follows: : 0=no infection; 1=maximum of one lesion per root; 2=more than one lesion per root; 3=many small and at least one large lesion per root; 4=many large lesions per root; 5=roots completely brown; 6=roots completely brown plus infection in stem; 7=roots completely brown, infection in stem and wilted yellow leaves; 8=entire plant brown and wilted (Fig. S20). The wheat plants that had germinated and grown in the absence of Take-all (with or without N2 spores) were healthy and infection free (Figs. S21 & S22). However, those seeds that had grown from un-inoculated seeds in the presence of the Take-all fungus showed extensive and severe levels of Take-all disease, with an average infection score of 7.24 ± 0.26 SE (Fig. S21). Most of the plants in this treatment group exhibited infected roots, stems and leaves, which all appeared senescent and brown (Fig. S22). However, there was a significant effect of plant treatment on infection score (Kruskal-Wallis test H_DF=3_= 83.41, *P* = <0.001). Plants that had grown from seeds coated in N2 spores (N = 25), demonstrated a small, but significant, decrease in average infection severity to 5.47 ± 0.59, compared to plants grown from uninoculated seeds in the presence of Take-all (Fig. S22, Dunn’s test between inoculated and sterile wheat grown with *G. graminis*: *P*= 0.023). As plants were grown in a sterile system in this experiment, it is likely that the streptomycete experienced low levels of nutrient availability compared to in a natural soil environment. We hypothesise that greater levels of competition and nutrients may fuel greater levels of antibiotic production by N2, and thus the strain could offer greater levels of protection against host infection in a more natural soil-plant system.

## Discussion

*Streptomyces* species have traditionally been described as free-living soil bacteria but given that most bare soils are rapidly colonised by vegetation it is perhaps not surprising that they are also effective at colonising the rhizosphere and endosphere of plants (10). It has even been suggested that their filamentous growth may have evolved to facilitate plant root colonisation since this trait evolved 50 million years after plants first colonised land, around 450mya (2, 10). Certainly, their ability to sporulate provides a useful mechanism for vertical transmission across plant generations, via the soil. In this study we followed up reports that *Streptomyces* bacteria are enriched in the *A. thaliana* rhizosphere and specifically recruited by plant-produced compounds such as the plant hormones salicylate and jasmonate (12–17). We aimed to isolate and characterise plant-associated streptomycetes from *A. thaliana* and test whether they can be beneficial to their plant host. We generated high quality genome sequences for five *A. thaliana* root-associated strains and three strains of the known endophyte *S. lydicus*. We found that three of the five root-associated strains significantly increased the biomass of *A. thaliana* plants when they were applied to seeds or roots, both *in vitro* and when applied in combination in soil. Two others, N1 and N2, significantly decreased the biomass of *A. thaliana in vitro*, most likely because they make polyenes that bind to sterols, including the phytosterols found in plant cell walls (44–46), which probably had a negative effect on the plant in a sterile system. However, this effect was removed in a compost system with neither strain influencing plant growth. Our work suggests therefore, that while *Streptomyces* are consistently associated with *A. thaliana* roots and enriched in the root microbiome compared to the surrounding bulk soil, not all strains that are competitive in the rhizosphere and endosphere necessarily have a beneficial effect on host fitness. This is important from an ecological and applied perspective, the former because it helps us to better understand the microbial and host factors influencing plant microbiome assembly and the latter because tipping the balance in the plant’s favour, for example by applying beneficial species as seed coatings or soil additives that can competitively colonise roots, could improve crop yields (19, 59, 60). Such strains could be used as biological growth promoters to replace the use of harmful pesticides and fertilisers which have negative effects on the wider ecosystem and also contribute to climate change (10, 11, 19, 59, 60). As a proof of this concept we took strain N2, which makes the polyene antifungals including pentamycin, 14-hydroxyisochainin and filipin III, and coated seeds of spring bread wheat with its spores. N2 was able to protect germinating wheat seedlings against the Take-all fungus *in vitro* and significantly reduce Take-all disease progression in wheat plants grown in sterile vermiculite. Although we do not have an assay for Take-all disease in soil the protective effect may be even greater because nutrients in the form of organic matter could provide a more beneficial growing environment for the *Streptomyces* strain than sterile vermiculite (61) and the presence of a greater diversity of microbes may fuel antimicrobial production by this strain as a result of interference competition (62). It is also intriguing that IAA increased the antifungal activity of strain N2 since it provides compelling evidence that environmental signals can alter the expression of secondary metabolites, and probably as an extension also influence interspecies competition (47, 49). Although there is much future work to do to understand this phenomenon our results provide a system to begin to understand how secondary metabolites are regulated and used by microbes in nature and may also provide new tools for activating the 90% of BGCs that are silent in these bacteria. Plant-associated *Streptomyces* strains may yet provide us with a new generation of antimicrobials for the clinic and might also be harnessed to improve our food security. Understanding the ecology of these bacteria and their associated natural products will be crucial if we are to achieve these goals.

## Materials and Methods

### Isolation of root-associated *Streptomyces* strains

Buffers and media recipes are listed in Table S8. Strains, primers and plasmids are listed in Table S1. Wild-type *A. thaliana* Col-0 seeds were sterilised by washing in 70% (v/v) ethanol for 2 minutes, 20% (v/v) sodium hypochlorite for 2 minutes, then five times in sterile water. Individual seeds were sown into pots of sieved Levington’s F2 seed and modular compost, placed at 4°C for 24 hours, then grown for 4 weeks under a 12-hour photoperiod (12 hours light/12 hours dark) at 22°C. Plants were taken aseptically from pots and their roots tapped firmly to remove as much adhering soil as possible. Root material was placed into sterile PBS-S buffer for 30 minutes on a shaking platform and before being transferred into fresh PBS-S and washed for 30 minutes. Any remaining soil particles were removed with sterile tweezers. Cleaned roots were then transferred to 25ml of fresh PBS-S and placed in a sonicating water bath for 20 minutes to remove any residual material still attached to the root surface; this was to ensure that any remaining bacteria were either present in the endophytic compartment or were very firmly attached to the root surface (“the rhizoplane”) (12). The roots were crushed in sterile 10% (v/v) glycerol and serial dilutions were spread onto either soya flour mannitol (SFM) agar, starch casein agar, or minimal salts medium agar containing sodium citrate. Plates were incubated at 30°C for up to 14 days. Colonies resembling streptomycetes were re-streaked onto SFM agar and identified by 16S rRNA gene PCR amplification and sequencing with the universal primers PRK341F and MPRK806R. Sequencing was carried out by Eurofins Genomics, Germany. Three strains of *Streptomyces lydicus,* which are known to associate with plant roots, were also used in experiments; *S. lydicus* WYEC108 was isolated from the commercial biocontrol product Actinovate® and two more *S. lydicus* strains (ATCC25470 and ATCC31975) were obtained from the American Type Culture Collection. *Streptomyces* strains were maintained on SFM agar (N1, N2, M2, M3), Maltose/Yeast extract/Malt extract (MYM) agar with trace elements (L2) or ISP2 agar (*S. lydicus* strains). Strains were spore stocked as described previously (63).

### Genome sequencing and analysis

High quality genome sequences were obtained for *Streptomyces* strains N1, N2, M2, M3, and L2, as well as the three known strains of *Streptomyces lydicus* using PacBio RSII sequencing technology at the Earlham Institute, Norwich, UK, as described previously (25). The automated Multi-Locus Species Tree (autoMLST) server (28) was used to phylogenetically classify the *Streptomyces* strains L2, M2, M3, N1 and N2. BGCs were predicted using antiSMASH 5.0 (31) and genomes were annotated using RAST (64). Amino acid sequences were uploaded to the KEGG Automatic Annotation Server (KAAS) for functional annotation of genes and metabolic pathway mapping (65).

### Generating eGFP-labelled *Streptomyces* strains

Plasmid pIJ8660 containing a codon optimised eGFP gene under the control of the constitutive *ermE*\* promoter and the *aac* apramycin resistance marker (66) was conjugated into *Streptomyces* strains (63). Exconjugants were selected and maintained on SFM agar plates (Table S8) containing 50 μg ml^−1^ apramycin. For confocal microscopy, *A. thaliana* Col-0 seeds were germinated on MSk (Table S8) and grown vertically at 22 °C for 9 days under a 12-hour photoperiod. Seedlings were then transferred to MSk (1.5% agar, 0% sucrose, Table S5) and allowed to equilibrate for 24 hours before being inoculated with 1μl of spore suspension (10^6^ spores ml^−1^) of the eGFP-tagged M3 or *Streptomyces coelicolor* M145 strains. As a known coloniser of plant roots, *S. coelicolor* was used as a control (32). Inoculated seedlings were then left to grow for 3 days before being washed in a 20% (v/v) solution of glycerol containing 1 μg ml^−1^ 276 SynaptoRed™ for 10 minutes. A 20 mm section of root (taken from the base of the petiole) was then mounted onto a slide with 100 μl of the SynaptoRed™/glycerol solution. Samples were imaged using a Zeiss LSM510 META laser-scanning confocal microscope with a PlanApochromat 63x (1.4 NA) objective. Green fluorescent protein was excited at 488 nm and emission collected through a 527.5 ± 22.5 nm bp filter, and FM4-64 excited at 543 nm and emission collected through a 587.5 ± 27.5 nm bp filter.

### Plant growth promotion assays

*A. thaliana* Col-0 seeds were sterilised and plated onto MSk medium (1% (w/v) sucrose and 0.8% (w/v) agar, Table S8). These were then left at 4 °C in the dark for 24 hours before being placed, vertically, under long day growth conditions (12 h light/ 12 h dark) at 22°C for 10 days. Seedlings were then transferred to square agar plates containing MSk agar (as above, Table S8) and allowed to equilibrate, vertically, overnight at 22 °C. 1 μl of *Streptomyces* spores (10^6^ ml^−1^) from each of the sequenced isolates was added to the top of the root system of each seedling and allowed to dry. 16 replicate seedlings were inoculated per sequenced *Streptomyces* strain. 10% (v/v) glycerol was added to control seedlings. Plates were grown vertically for 16 days, 12 h light/ 12 h dark at 22°C before measuring plant biomass (dry weight). The biomass of plants with different inocula were compared via ANOVA and Tukey’s Honestly Significant Difference (HSD) tests using R 3.2.3 (67); biomass was log-transformed during analyses to ensure normality of residuals. Strains were also tested for their ability to promote *A. thaliana* growth in compost. Sterile *A. thaliana* Col-0 seeds were placed into a solution of 2xYT (Table S8) containing 10^6^ pregerminated spores ml^−1^ of each strain or no spores as a control. Seeds were incubated in the spore solution for 2 hours, before being transferred to pots containing sieved Levington’s F2 seed and modular compost. An additional 3 ml of pre-germinated spores (or sterile 2xYT) was pipetted into the soil surrounding each seed. The strains L2, M2 and M3 were also tested for their ability to promote plant growth in combination; 10^3^ spores ml^−1^ of each strain were mixed together and pregerminated in 2xYT before being used as above. Pots were then placed at 4°C for 48 hours before being grown for 6 weeks under a photoperiod of 12 hours light/12 hours dark. There were 8 replicate pots per treatment. After 6 weeks, the plants were removed from pots and cleaned by washing in PBS-S (Table S8) and using tweezers to remove adhering soil particles. Plants were then dried in an oven at 50°C enabling plant dry weight to be calculated. An ANOVA test and Tukey’s HSD tests were used (as above) to test for an effect of strain inoculation on plant dry weight. Dry weights were log-transformed prior to analysis.

### Indole 3 Acetic Acid (IAA) production assays

*Streptomyces* isolates were grown on cellophane membranes covering YMD media supplemented with 5 mM tryptophan (Table S8). After 7 days, cellophane membranes with bacterial biomass were removed and plates were flooded with Salkowski reagent (as described in 36). A red colour indicates that IAA has leached into the medium.

### 1-Aminocyclopropane-1-carboxylic acid(ACC) degradation assays

To test for the use of ACC as a sole nitrogen source, *Streptomyces* strains were streaked onto Dworkin and Foster medium (68) in which 0.2% (w/v) NH_4_SO_4_ or 0.051% (w/v) ACC was added as a sole nitrogen source, or no nitrogen source as a control. Plates were incubated for 10 days at 30°C before imaging.

### Antibiotic bioassays

Spores (4 μl of 10^6^ ml^−1^ solution) of individual *Streptomyces* isolates were pipetted onto the centre of agar plates and incubated at 30°C for 7 days before adding the pathogenic indicator strains (see Table S1). A clinical isolate of *Candida albicans* (gift from Prof Neil Gow), *Bacillus subtilis* 168 (from Prof Nicola Stanley Wall), a clinical isolate of Methicillin Resistant *Staphylococcus aureus* isolated from a patient at the Norfolk and Norwich University Hospital UK (69), an *Escherichia coli* K12 lab strain and the plant pathogen *Pseudomonas syringae* DC3000 (gift from Dr Jacob Malone) were grown overnight in 10 ml Lysogeny Broth (LB) at 30°C, 250 rpm. These were sub-cultured 1 in 20 (v/v) for a further 4 hours at 30°C before being used to inoculate 100 ml of molten LB (0.5% agar), 3 ml of which was used to overlay each agar plate containing a *Streptomyces* colony. Plates were incubated for 48 hours at 30°C. Bioactivity was indicated by a clear halo around the *Streptomyces* colony. For bioassays using the fungal strains *Lomentospora prolificans* or *Gaeumannomyces graminis* var. *tritici* (Table S1), a plug of the fungus (grown for 14 days on potato glucose agar) was placed at the edge of the agar plate, 2 cm away from the growing streptomycete colony. Plates were incubated at 25°C for up to 14 days to assess inhibition of fungal growth. Bioassays were carried out on a range of different media, including minimal medium supplemented with Indole-3-Acetic Acid (IAA) (See Table S8 for media recipes).

### Purification and elucidation of filipin-like compounds from strain N2

Spores of strain N2 were spread onto 120 plates (4 L) of SFM agar and grown for eight days at 30 °C. The resulting agar was then sliced into small pieces and extracted with ethyl acetate (3 × 1.5 L). An analytical sample was taken for analysis whereby the extract was filtered through gauze and the solvent removed under reduced pressure yielding 9.2 g of crude material. This was split into two halves and each treated identically: after dissolving in acetone (50 ml), loose normal phase silica gel (~30 g, Sigma Aldrich) was added and the solvent removed under reduced pressure. The impregnated silica gel was dry loaded onto a Biotage® SNAP Ultra cartridge (50 g, HP-Sphere normal phase silica). The resulting sample was chromatographed using a Biotage flash chromatography system to separate the polyene fraction (338 nm) and the actinomycin(s) containing fraction (444 nm; hereafter referred to as the ‘actinomycin complex’) using the following gradient with a flow rate of 100 ml min^−1^: hold at 0% B for 1 CV; linear gradient 0-50% B over 10 CV; 50-100% B over 0.5 CV; and hold at 100 % B for 3 CV; (mobile phase A, chloroform; mobile phase B, methanol).

The polyene containing fractions were combined, the solvent removed under reduced pressure and the residues split into two fractions. Each fraction (in 800 μL DMSO) was loaded onto a Biotage® SNAP Ultra cartridge (12 g, C_18_) and chromatography was achieved using the following gradient at a flow rate of 12 ml min^−1^: hold at 0% B for 5 CV; linear gradient 0-55% B over 1 CV, 55-85% B over 10 CV; 85-100% B over 2 CV; and hold at 100% B for 1 CV; (mobile phase A, water; mobile phase B, methanol). The resulting fractions were analysed using LCMS and combined to yield three polyene samples of 53 mg, 34 mg and 9 mg after solvent was removed. LCMS spectra of these fractions and the original crude extract were then uploaded onto the GNPS (Global Natural Products Social Molecular Networking platform). The largest network containing the spectra of the filipin related compounds was manually adapted and processed in Cytoscape 3.6.1. The second fraction contained the most interesting compounds so only this sample was further purified by chromatography over a Synergi Fusion 4 micron C_18_ 250 × 10 mm column (Phenomenex) using an Agilent 1100 series HPLC system fitted with a fraction collector and eluting at a flow rate of 3.5 ml min^−1^ with the following gradient. 0-2 min, 45% B; 2-5 min, 45-50% B; 5-10 min, 50% B; 10.0-10.1 min, 50-45% B; 10.1-12.0 min, 45% B; (mobile phase A, 0.1% formic acid in water; mobile phase B, acetonitrile). This yielded pentamycin (4.2 mg) and 14-hydroxyisochainin (2.3 mg). Both structures were assigned using 2D NMR recorded on a Bruker Avance Neo 600 MHz spectrometer equipped with a helium-cooled cryoprobe and dissolved in DMSO-*d*_6_. Additional 1D-experiments were carried out in CD_3_OD for 14-hydroxyisochainin in order to compare directly with published data for this compound (Figs S10-16 and Tables S5-8) (52).

Disc-diffusion bioassays were used to test whether purified pentamycin, 14-hydroxyisochainin and the actinomycin complex were active against the pathogenic strains *C. albicans* and *E. coli* (Table S1). Both indicator strains were grown in 10 ml of LB broth (Table S5) at 30°C and 200 rpm overnight. Cultures were then diluted 1 in 20 (v/v) into 10 ml of LB and grown for a further 4 hours. The 10 ml sub-culture was added to 50 ml of soft LB (0.5% agar) which was then poured into 10 cm square plates and allowed to set. Meanwhile, 6 mm sterile filter paper discs (Whatman) were inoculated with 40 μl of each individual compound (three technical replicates of each compound were tested). 40 μl of methanol was added to one disc per plate as a solvent control and 40 μl of nystatin (5 mg ml^−1^) or hygromycin (50 mg ml^−1^) were used as positive controls on *C. albicans* and *E. coli* plates, respectively. Once dry, discs were placed onto plates 3 cm apart. These were then incubated at 30°C overnight. Inhibition of the indicator strain was evidenced by a zone of clearing around the disc. Purified compounds were also tested for their ability to inhibit the wheat take-all fungus *G. graminis* (Table S1). For this, discs were placed onto PGA agar plates (Table S8) 2 cm away from an actively growing plug of *G. graminis*. Plates were left to grow at room temperature for 5 days before imaging. Three technical replicates were run for each purified compound (pentamycin, 14-hydroxyisochainin and actinomycin).

### Wheat seedling bioassays with *Streptomyces* strain N2

Seeds of *Triticum aestivum* (var. Paragon, Table S1) were sterilized by placing them in 70% (v/v) ethanol for 2 minutes followed by a wash in 3% (v/v) NaOCl for 10 minutes. Seeds were then rinsed five times in sterile dH_2_O, before placing them into a solution of pregerminated spores (10^7^ spores ml^−1^) of *Streptomyces* strain N2 (Table S1). Spores were pregerminated in 2xYT (Table S8) at 50°C for 10 minutes. Seeds were incubated in either the spore preparation or sterile 2xYT as a control, for 2 hours, before being allowed to dry in a Petri dish under sterile conditions. Two wheat seeds were then placed onto a 10 cm square plate of Msk agar (1.5% (w/v) agar, 0% (w/v) sucrose, Table S8), on either side of a plug of the *G. graminis* fungus, which was placed in the centre of the agar plate. Plugs were taken from an actively growing plate of *G. graminis* on PGA agar. Three replicate plates each of N2-coated seeds and sterile control seeds were used in each experiment. Plates were incubated for 5 days at room temperature after which inhibition of *G. graminis* was indicated by a zone of clearing around the wheat seeds.

A sterile vermiculite system was used to investigate the ability of *Streptomyces* strain N2 to protect older wheat seedlings against Take-all infection. 25 ml of sterile vermiculite was placed into the bottom of a 50 ml Falcon tube. Five plugs of *G. graminis* actively growing on PGA (Table S8), or uninoculated PGA plugs as a control, were placed on top of this layer, before being covered with a further 10 ml of vermiculite. Five seeds of *T. aestivum* (soaked in either N2 spores or uninoculated 2xYT, as described above) were then placed on top of this vermiculite layer, before the addition of a further 10 ml of vermiculite. The Falcon tubes were then sealed with parafilm and incubated at 25°C for 3 weeks, under a 12 hour light/ 12 hour dark photoperiod. There were five replicates tubes, each containing five replicate seeds, of each of the following combinations: PGA plugs with N2-coated seeds (wheat/*Streptomyces* control); *G. graminis* plugs with N2-coated seeds (wheat/*Streptomyces*/fungus treatment); PGA plugs with uninoculated seeds (wheat control); *G. graminis* plugs with uninoculated seeds (wheat/fungus treatment). After three weeks of incubation, plants were taken from the Falcon tubes and adhering vermiculite was removed from the roots. Take-all infection severity was scored on a scale of 0-8, using an infection scoring system as follows: : 0=no infection; 1=maximum of one lesion per root; 2=more than one lesion per root; 3=many small and at least one large lesion per root; 4=many large lesions per root; 5=roots completely brown; 6=roots completely brown plus infection in stem; 7=roots completely brown, infection in stem and wilted yellow leaves; 8=entire plant brown and wilted. Differences in infection score between treatments were analysed in R 3.2.3 (67) using a Kruskal-Wallis test, coupled with a post-hoc Dunn’s multiple comparison test.

## Acknowledgments

SFW was funded by a Natural Environment Research Council (NERC) PhD studentship (NERC Doctoral Training Programme grant NE/L002582/1), JN was supported by a Biotechnology and Biological Sciences Research Council (BBSRC) PhD studentship (BBSRC Doctoral Training Programme grant BB/M011216/1), JR was funded by Leonardo Office Thuringia within the scope of Erasmus+ and the Deutschland stipendium, NAH was funded by NERC responsive mode grant NE/M015033/1 awarded to MIH, BW and JCM, and SFDB was funded by BBSRC responsive mode grant BB/P021506/1 to BW. Confocal microscopy was carried out at the Henry Wellcome Laboratory for Cell Imaging at UEA and we thank Dr Paul Thomas for training JN. This work was also supported by the Norwich Research Park (NRP) through a Science Links Seed Corn grant to MIH and JCM and by the NRP Earth and Life Systems Alliance (ELSA). We thank Anne Osbourn and Rachel Melton at the John Innes Centre (JIC) for assistance with the Take-all infection assays, Jacob Malone (JIC) for gifting us the *Pseudomonas* strain and Simon Orford (JIC) for the Paragon wheat seeds. Media recipes and other *Streptomyces* resources are available at ActinoBase.org.

