## Supplementary Figures and Tables for "*Streptomyces* endophytes promote the growth of *Arabidopsis thaliana*"

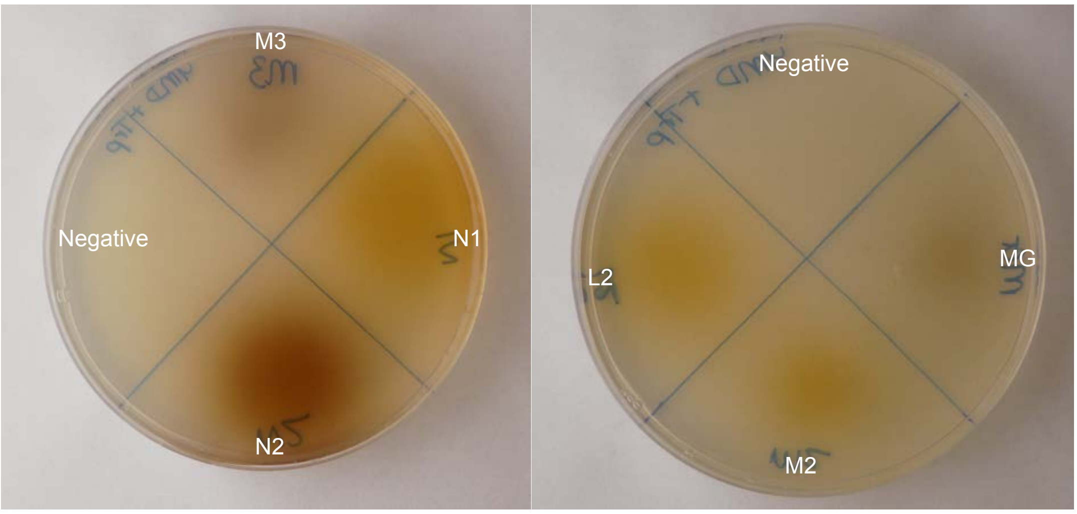

**Figure S1.** The *in vitro* production of IAA by *Streptomyces* isolates M3, N1, L2, N2, M2 and MG. Isolates were grown on cellophanes covering YMD agar supplemented with 5 mM tryptophan for 7 days. Cellophanes were removed and plates were flooded with Salkowski reagent. A red/pink colour indicates IAA has leached into the media. Negative = no bacteria were grown on this part of the cellophane.

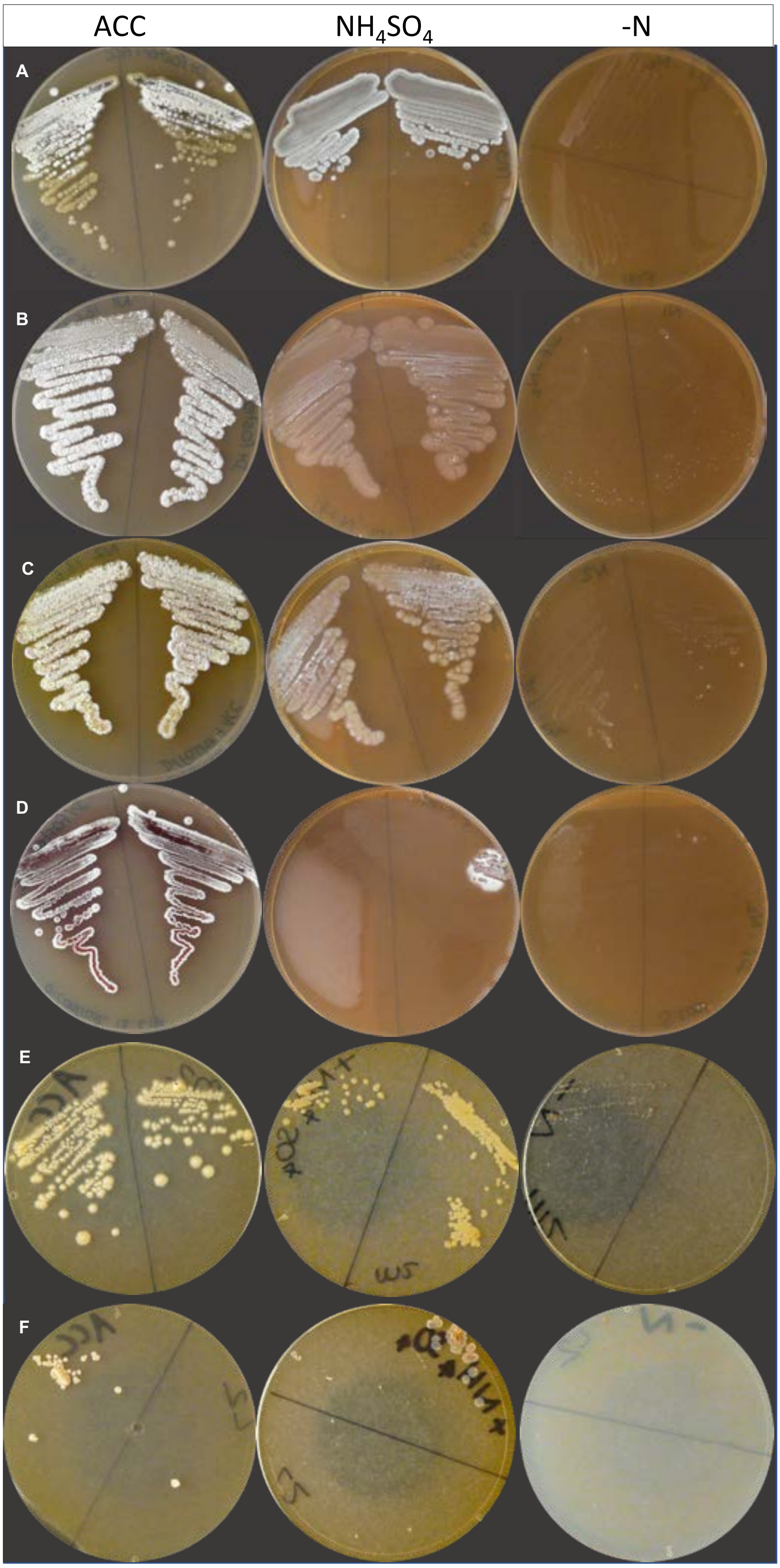

**Figure S2.** The use of ACC as a sole nitrogen source. *Streptomyces* strains A) M3, B) N1, C) N2, D) *Streptomyces coelicolor* M145, E) M2 and F) L2 were grown on Dworkin and Foster medium containing either 1-aminocyclopropane-1-carboxylic acid (ACC) or $NH_4(SO)_4$ as a sole nitrogen source, or no nitrogen (-N) as a control.

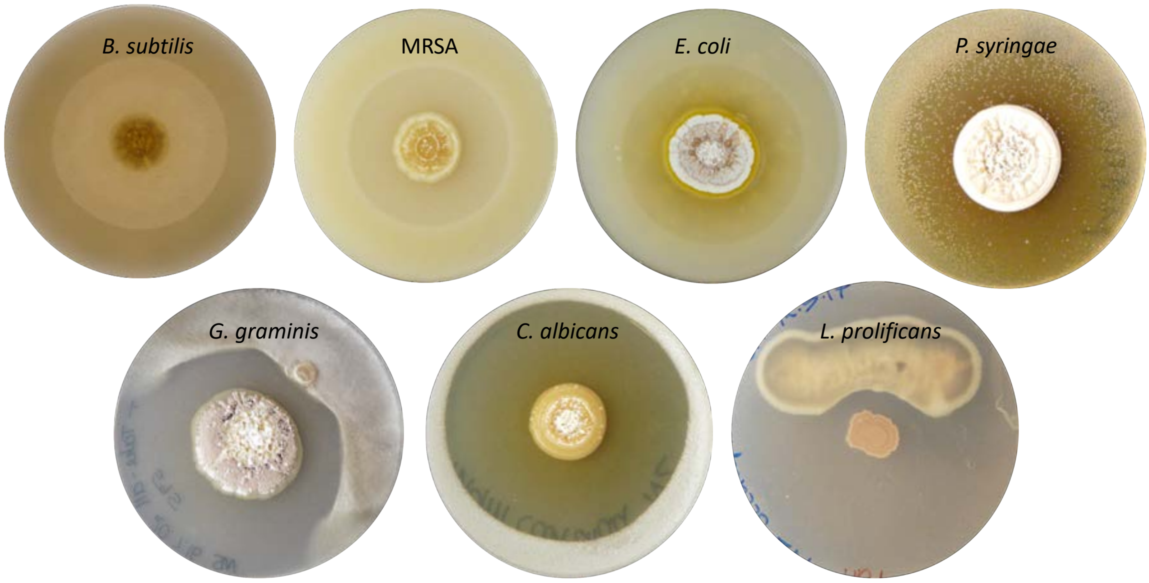

**Figure S3.** Bioassay plates demonstrating the ability of the *Streptomyces* isolate N2 (inoculated on the centre of the plates) to inhibit the growth of the Gram-positive bacteria *B. subtilis* and methicillin-resistant *Staphylococcus aureus* (MRSA), the Gram-negative bacteria *E. coli* and *P. syringae* as well as the fungal pathogens *Candida albicans*, *G. graminis* var. *tritici* (Take-all fungus) and *Lomentospora prolificans*.

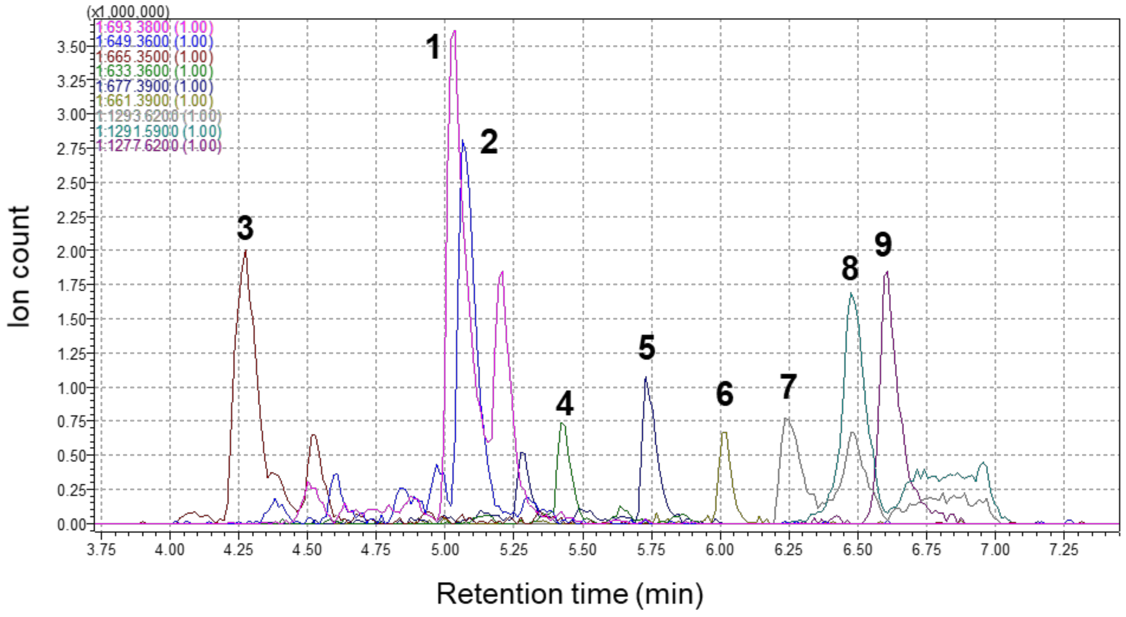

| Peak number | Measured mass $[M+Na]^+$ | Calculated mass $[M+Na]^+$ | Δppm | Molecular Formula | Compound assignment |
|---|---|---|---|---|---|
| 1 | 693.3842 | 670.3928 | 2.2 | $C_{35}H_{58}O_{12}$ | pentamycin |
| 2 | 649.3585 | 626.3666 | 2.7 | $C_{33}H_{54}O_{11}$ | 14-hydroxyisochainin |
| 3 | 665.3491 | 642.3615 | 1.6 | $C_{33}H_{54}O_{12}$ | 1´,14-dihydroxyisochainin |
| 4 | 633.3624 | 610.3717 | 1.5 | $C_{33}H_{54}O_{10}$ | isochainin |
| 5 | 677.3845 | 654.3979 | 2.6 | $C_{35}H_{58}O_{11}$ | filipin III |
| 6 | 661.3959 | 638.4030 | 3.7 | $C_{35}H_{58}O_{10}$ | filipin II |
| 7 | 1293.6184 | 1270.6234 | 5.8 | $C_{62}H_{86}N_{12}O_{17}$ | actinomycin $X_{0β}$ |
| 8 | 1291.5936 | 1268.6077 | 3.4 | $C_{62}H_{84}N_{12}O_{17}$ | actinomycin $X_2$ |
| 9 | 1277.6192 | 1254.6285 | 1.5 | $C_{62}H_{86}N_{12}O_{16}$ | actinomycin D |

**Figure S4.** Ion chromatograms from UPLC-MS analysis of the crude ethyl acetate extract from *Streptomyces* strain N2 grown on SFM agar plates. Numbers above the peaks on the chromatogram represent the different compounds identified in the extract. The main compounds present are summarized in the table beneath the chromatogram.

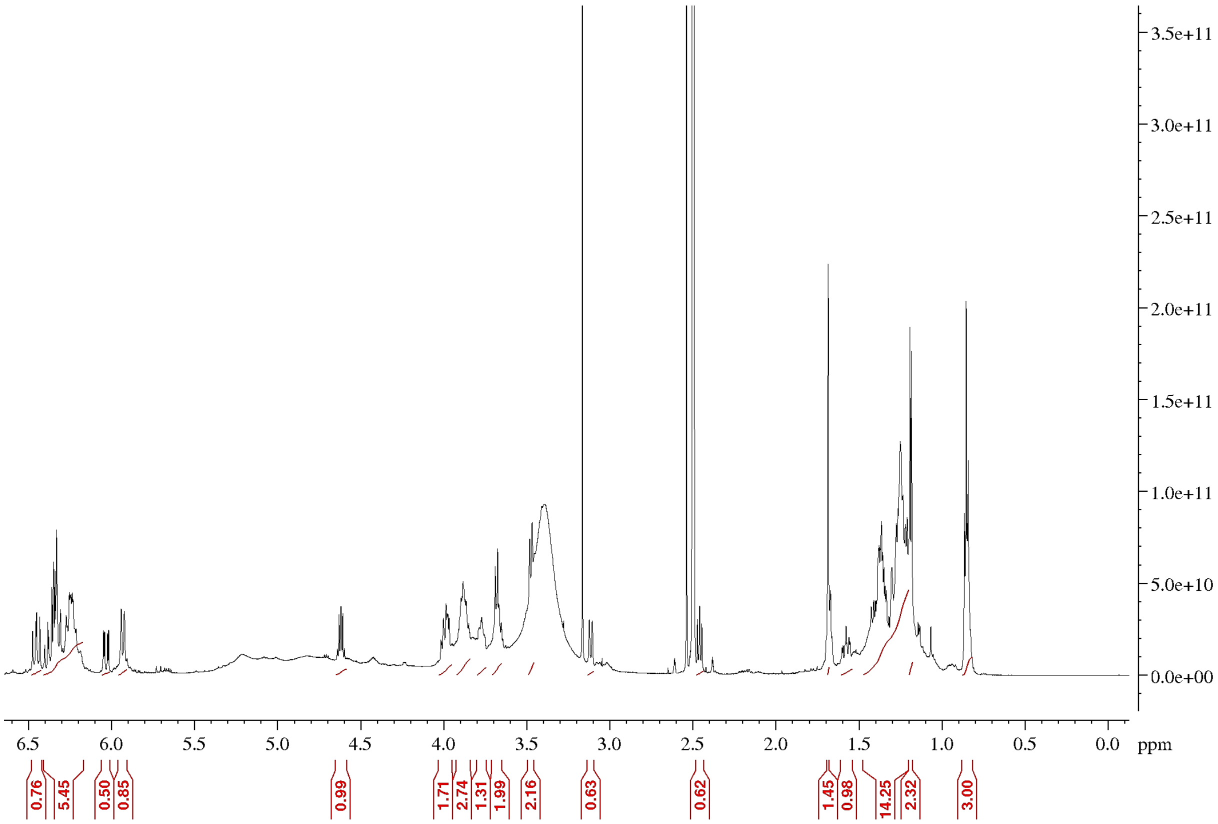

**Figure S5.** $^{1}$H NMR spectrum of pentamycin at 600 MHz in DMSO-$d_6$.

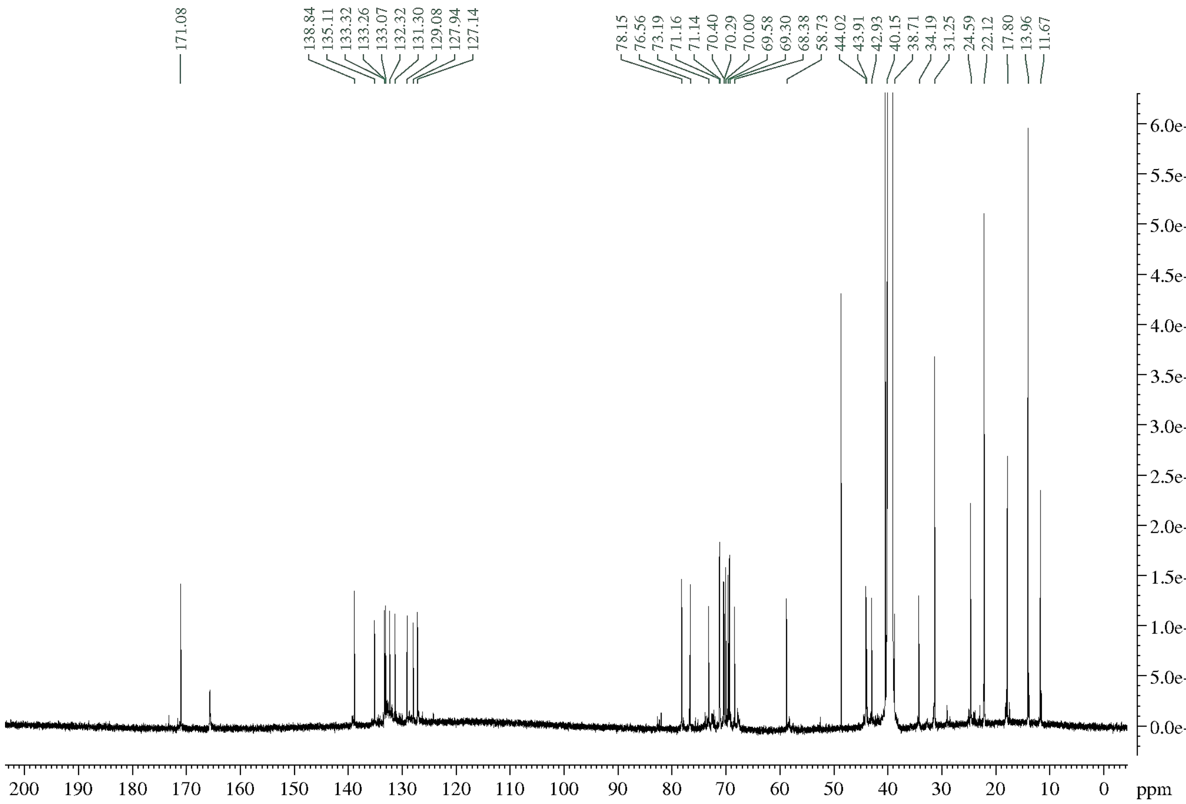

**Figure S6.** $^{13}C$ NMR spectrum of pentamycin at 150 MHz in DMSO-$d_6$.

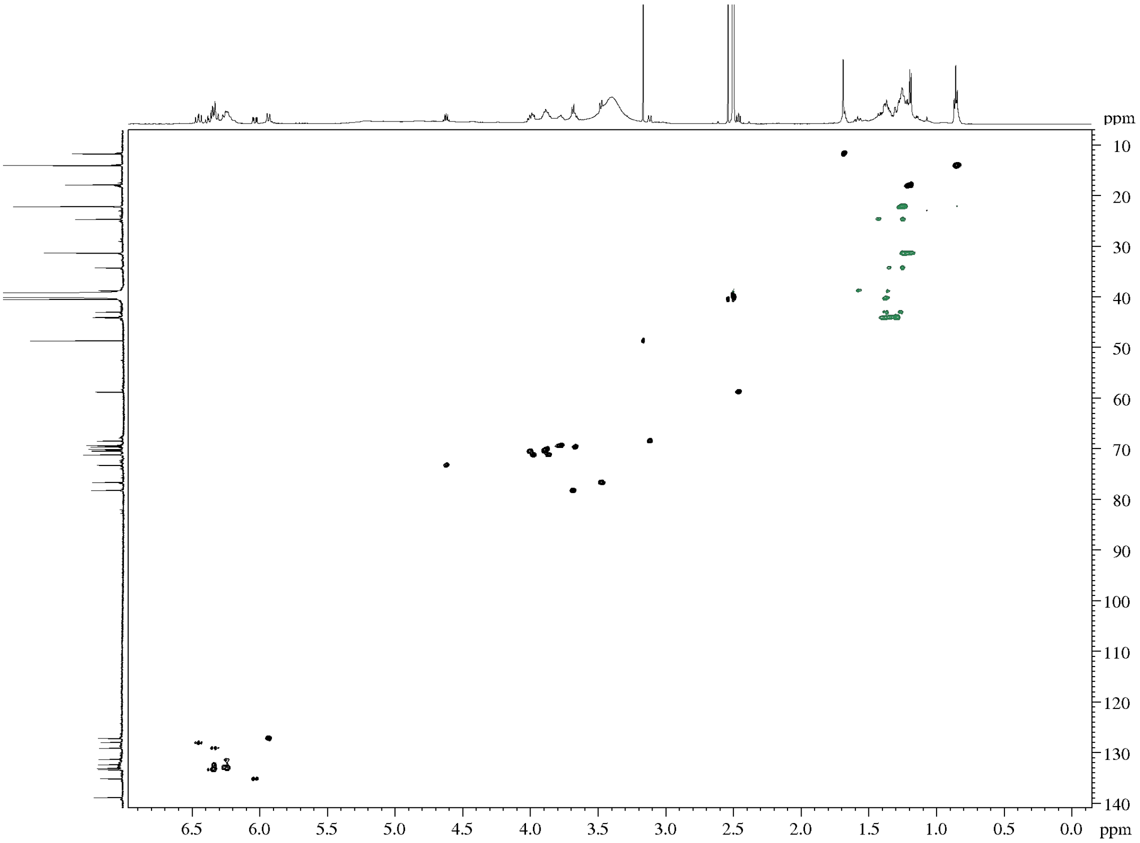

**Figure S7.** HSQCed spectrum of pentamycin in DMSO-$d_6$.

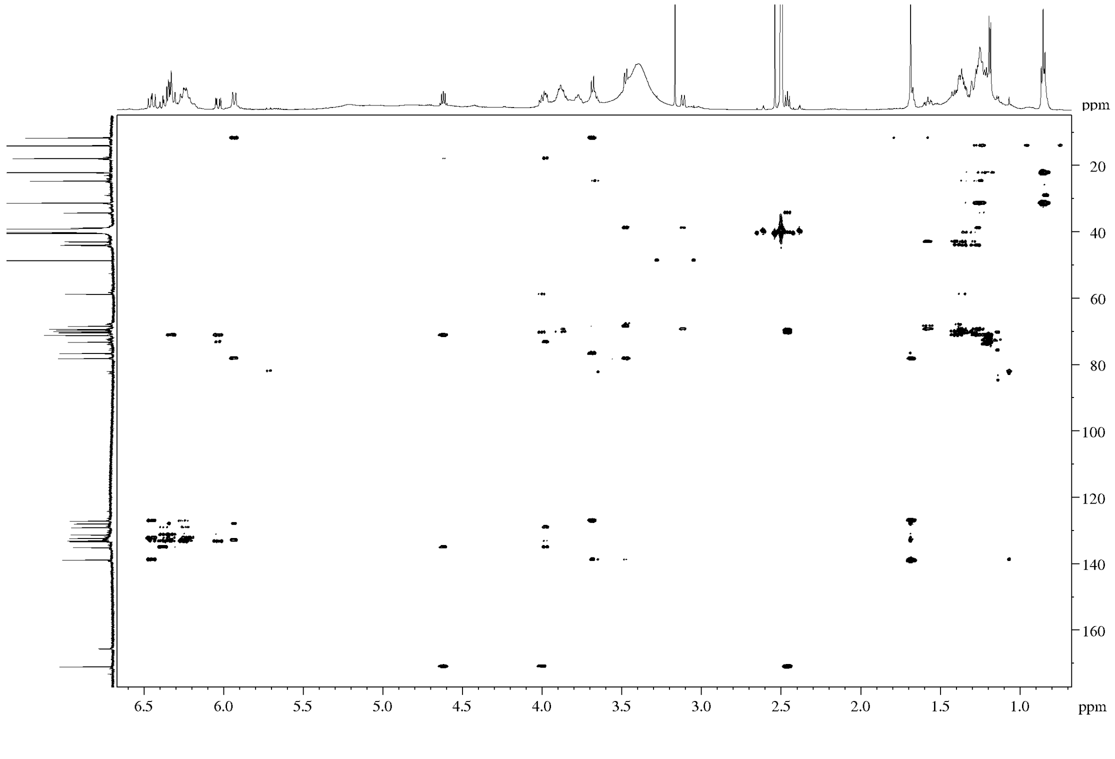

**Figure S8.** HMBC spectrum of pentamycin in DMSO-$d_6$

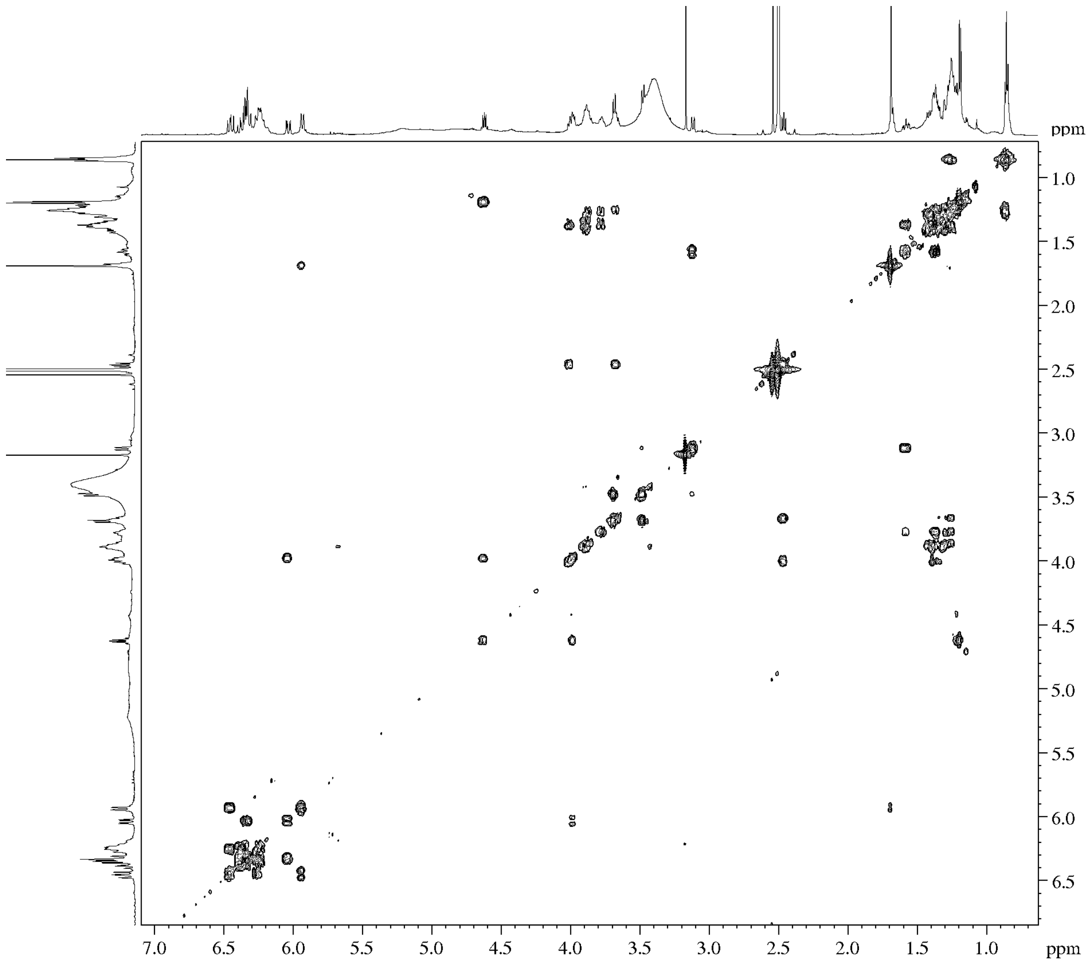

**Figure S9.** COSY spectrum of pentamycin in DMSO-$d_6$.

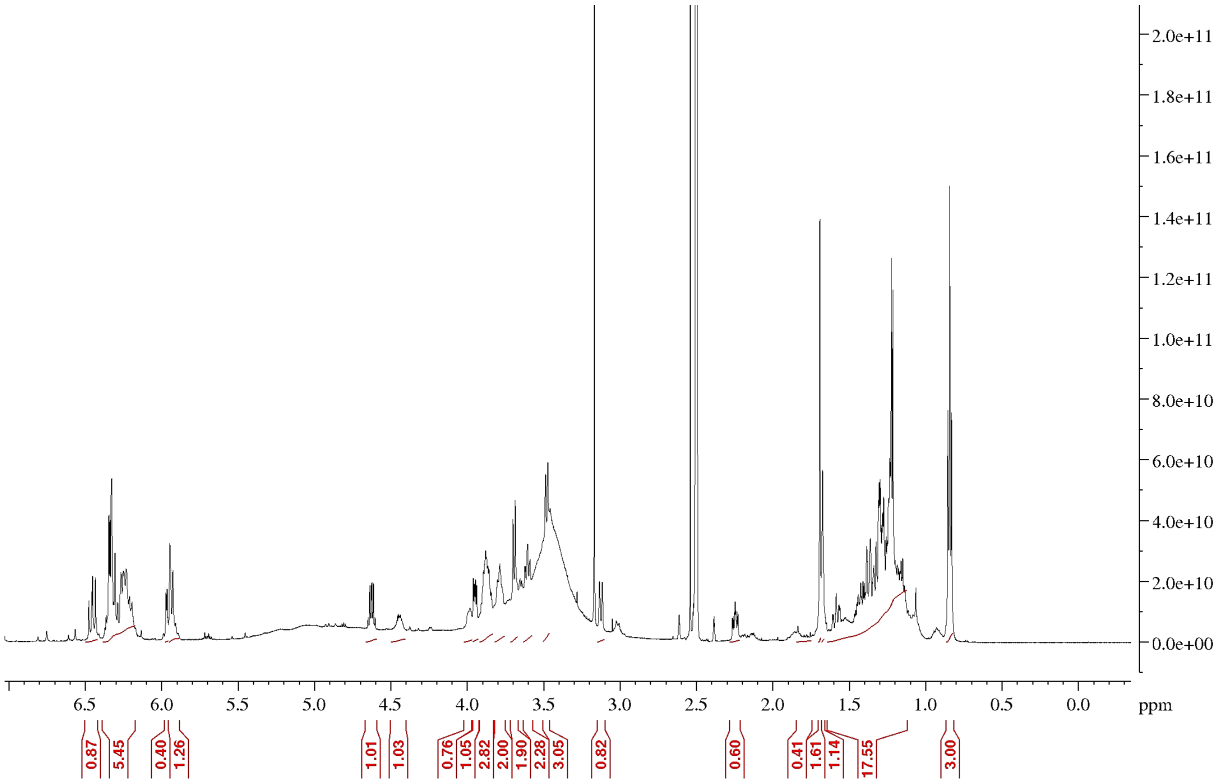

**Figure S10.** $^{1}$H NMR spectrum of 14-hydroxyisochainin at 600 MHz in DMSO-$d_6$.

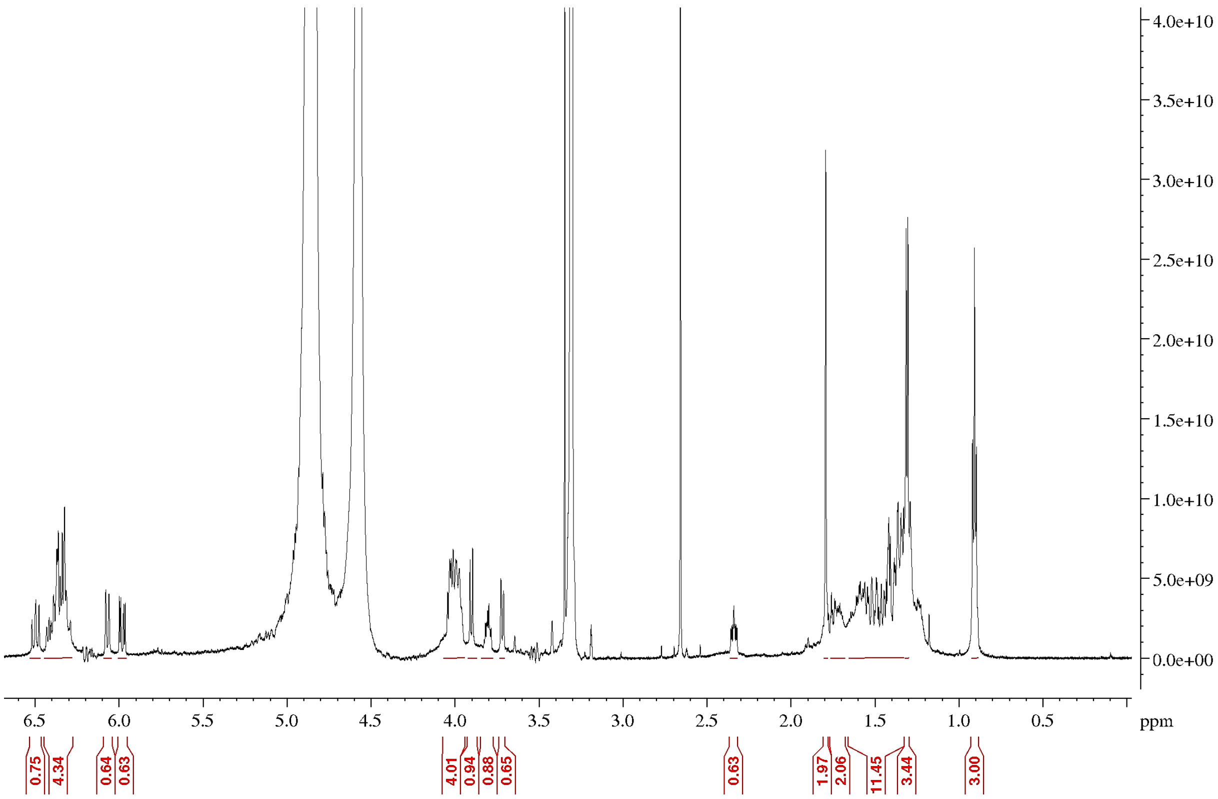

**Figure S11.** $^{1}$H NMR spectrum of 14-hydroxyisochainin at 600 MHz in $CD_3OD$.

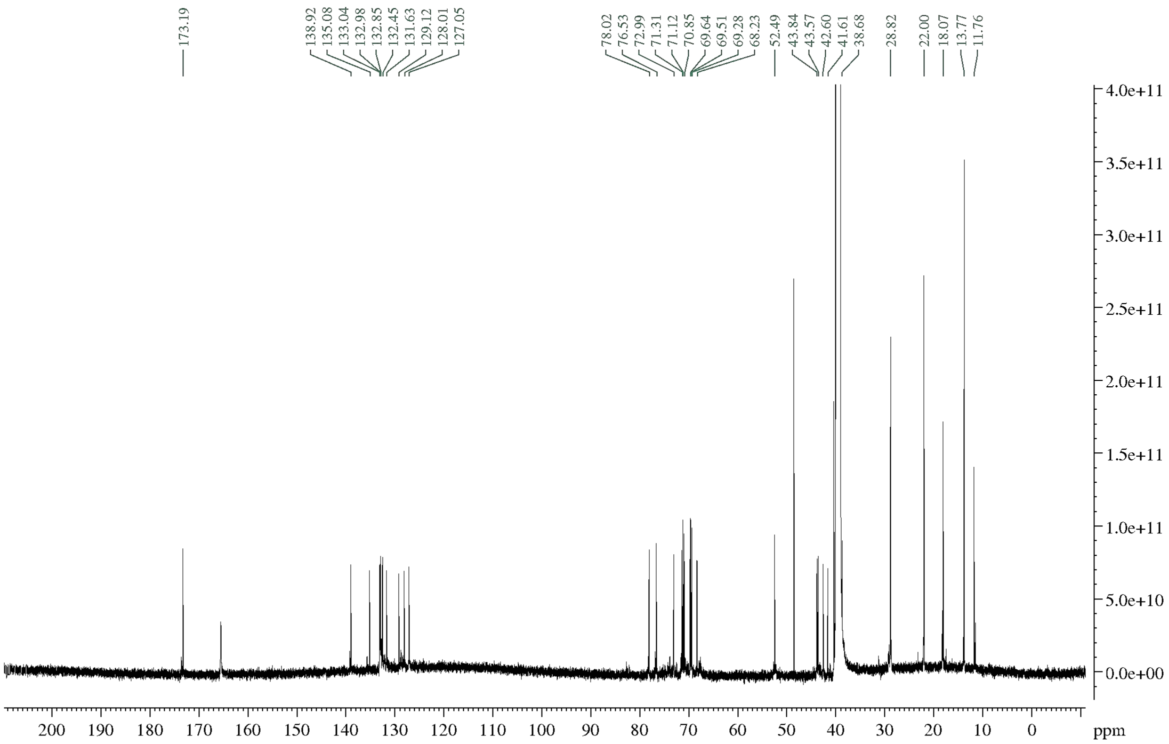

**Figure S12.** $^{13}$C NMR spectrum of 14-hydroxyisochainin at 150 MHz in DMSO-$d_6$.

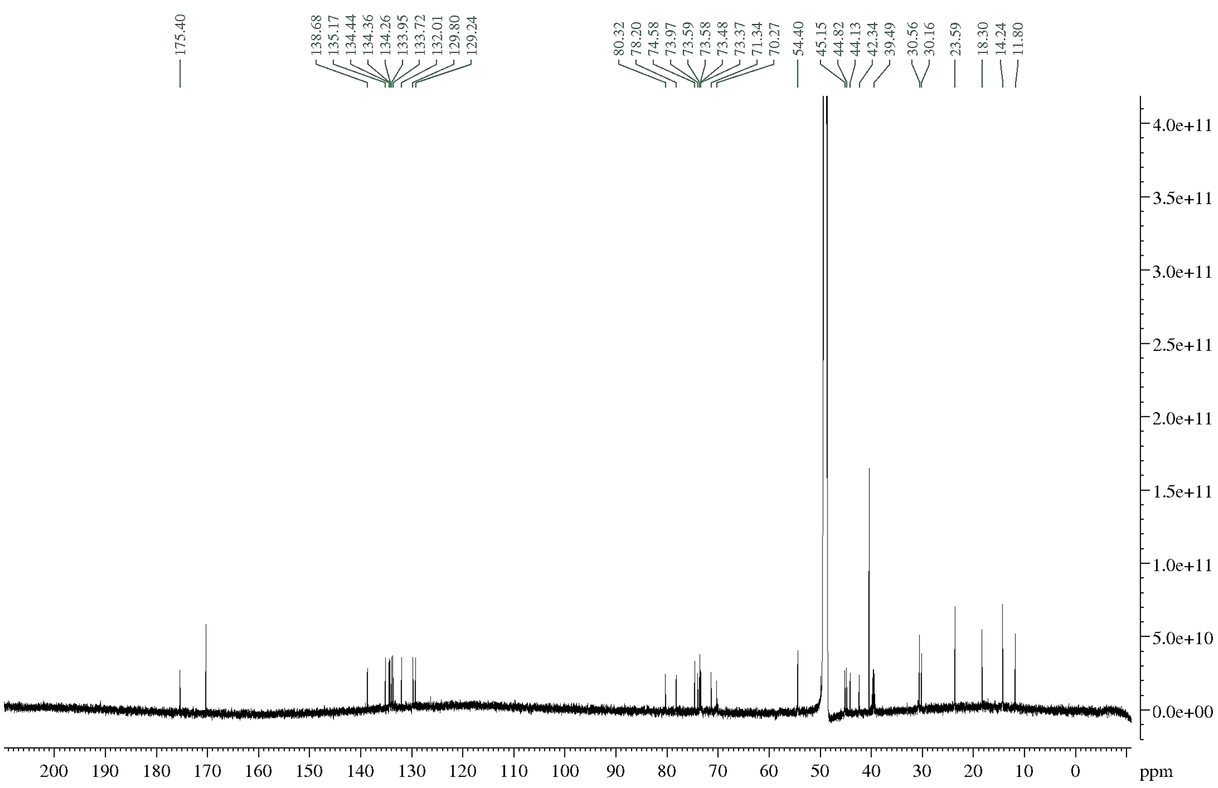

**Figure S13.** $^{13}$C NMR spectrum of 14-hydroxyisochainin at 150 MHz in $CD_3OD$.

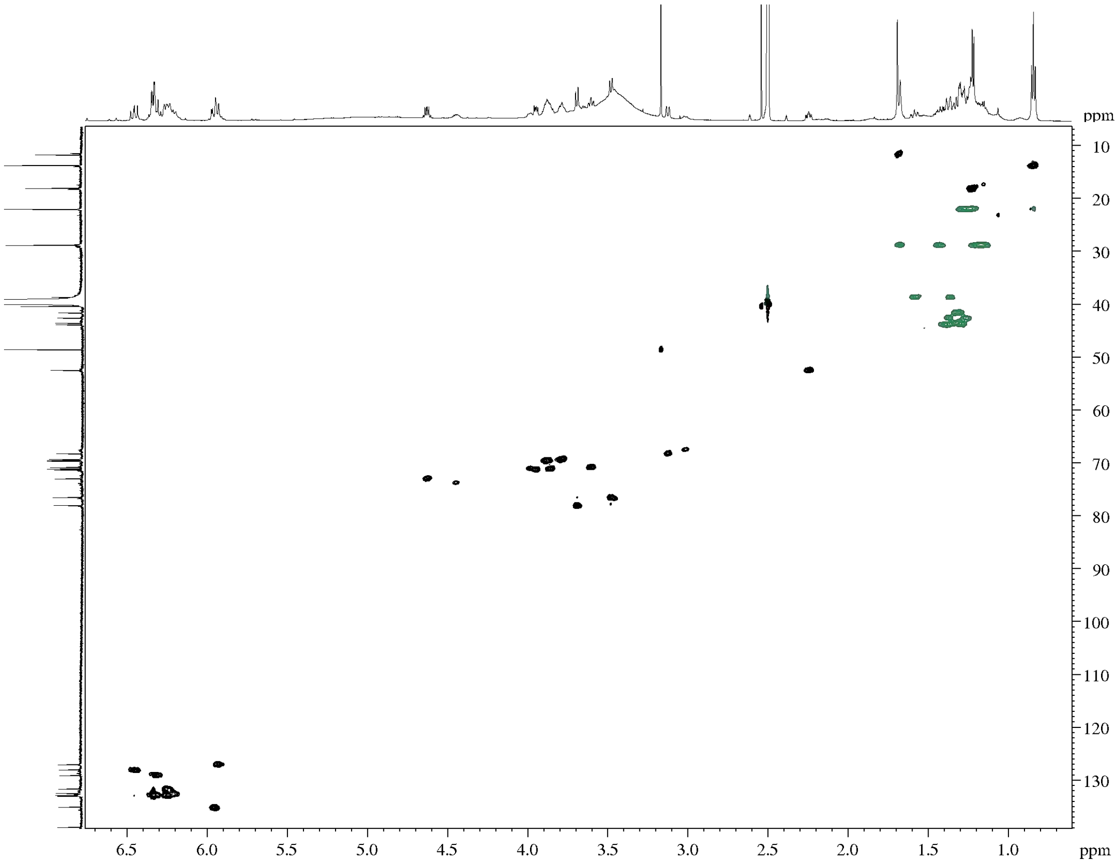

**Figure S14.** HSQCed spectrum of 14-hydroxyisochainin in DMSO-$d_6$.

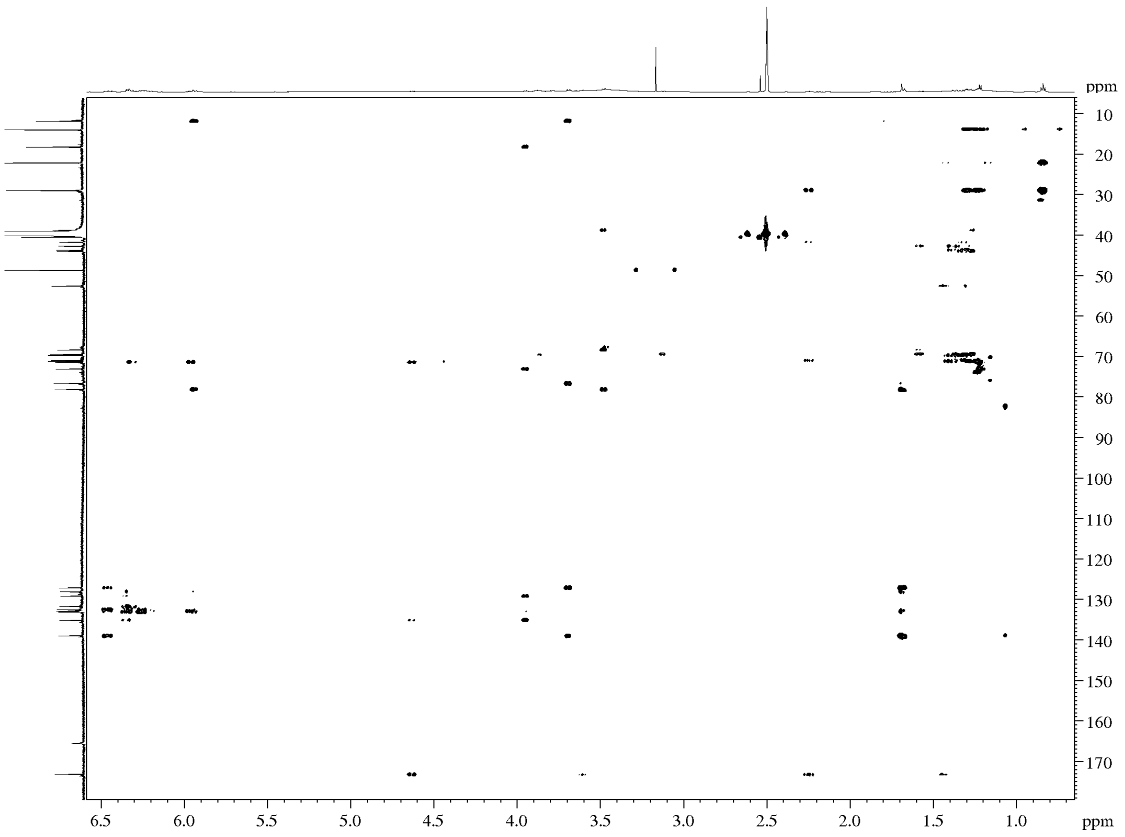

**Figure S15.** HMBC spectrum of 14-hydroxyisochainin in DMSO-$d_6$.

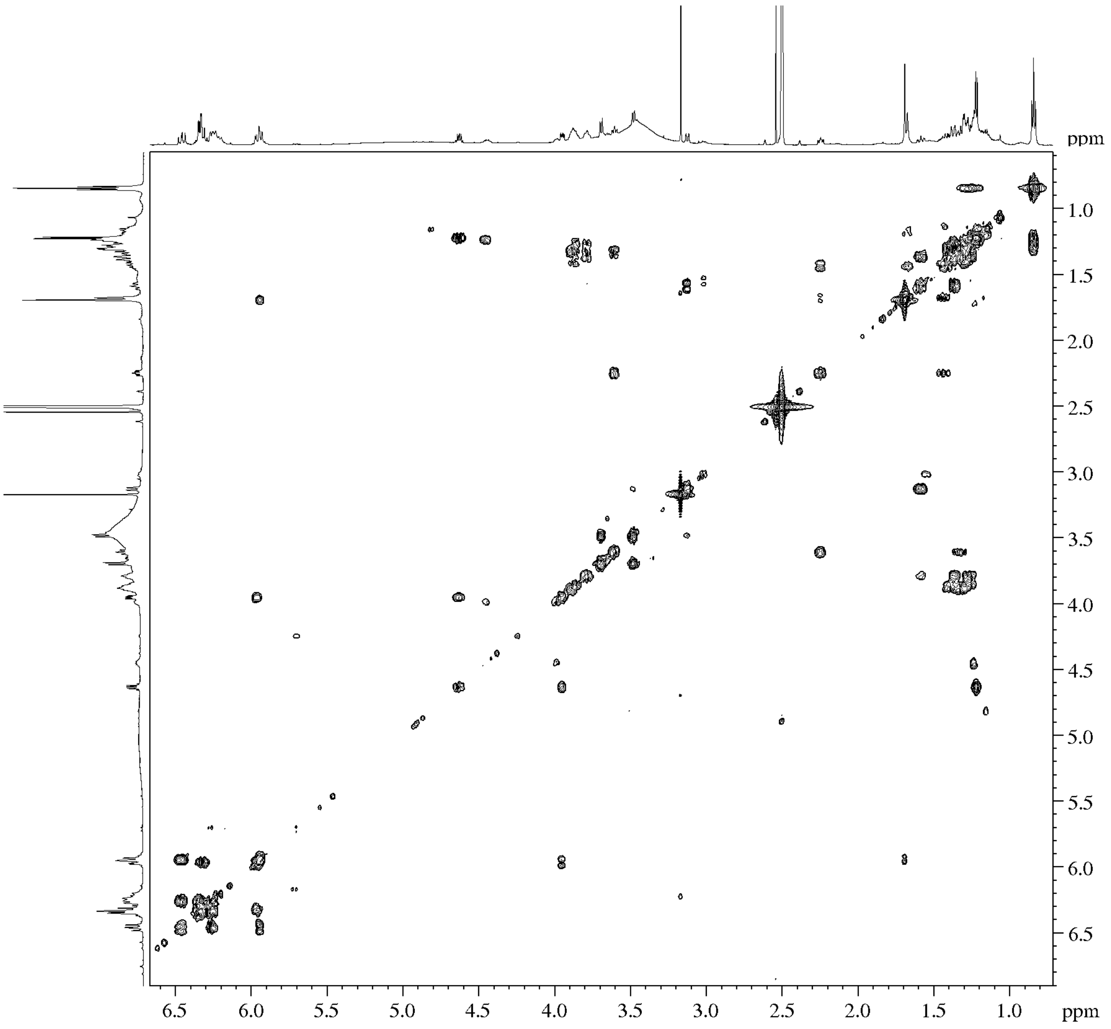

**Figure S16. COSY** spectrum of 14-hydroxyisochainin in DMSO-$d_6$.

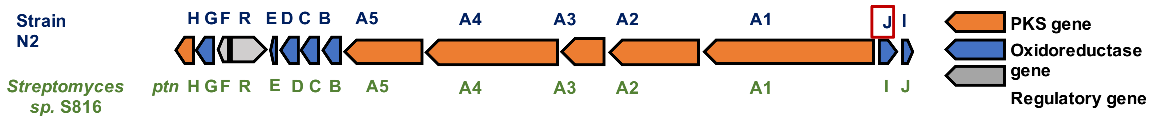

| N2 Protein | *Streptomyces sp.* S816 | Percentage identity (%) | Proposed function |
|---|---|---|---|
| I | ptnI | 98 | Ferredoxin for PtnJ |
| J | PtnJ | 100 | Cytochrome P450 hydroxylase |
| A1 | ptnA1 | 94 | PKS |
| A2 | ptnA2 | 95 | PKS |
| A3 | ptnA3 | 95 | PKS |
| A4 | ptnA4 | 96 | PKS |
| A5 | ptnA5 | 96 | PKS |
| B | ptnB | 93 | Putative dehydrogenase |
| C | ptnC | 99 | Cytochrome P450 hyrdoxylase |
| D | ptnD | 98 | Cytochrome P450 hyrdoxylase |
| E | ptnE | 94 | Ferrodoxin |
| R | ptnR | 91 | DnRI/RedD/AfsR-family transcriptional regulator |
| F | PtnF | 95 | LuxR family transcriptional regulator |
| G | PtnG | 99 | Putative oxidase |
| H | PtnH | 97 | Thioesterase |

**Figure S17.** A comparison of the architecture of the biosynthetic gene cluster encoding filipin-like compounds in *Streptomyces* N2 and *Streptomyces* sp. S816 (51), respectively. Filipin biosynthesis is encoded by 13 genes (A1-H) and pentamycin is formed as a result of the hydroxylation of filipin III by an additional cytochrome P450 monooxygenase (J, outlined in red), the gene for which is located upstream of the main filipin cluster.

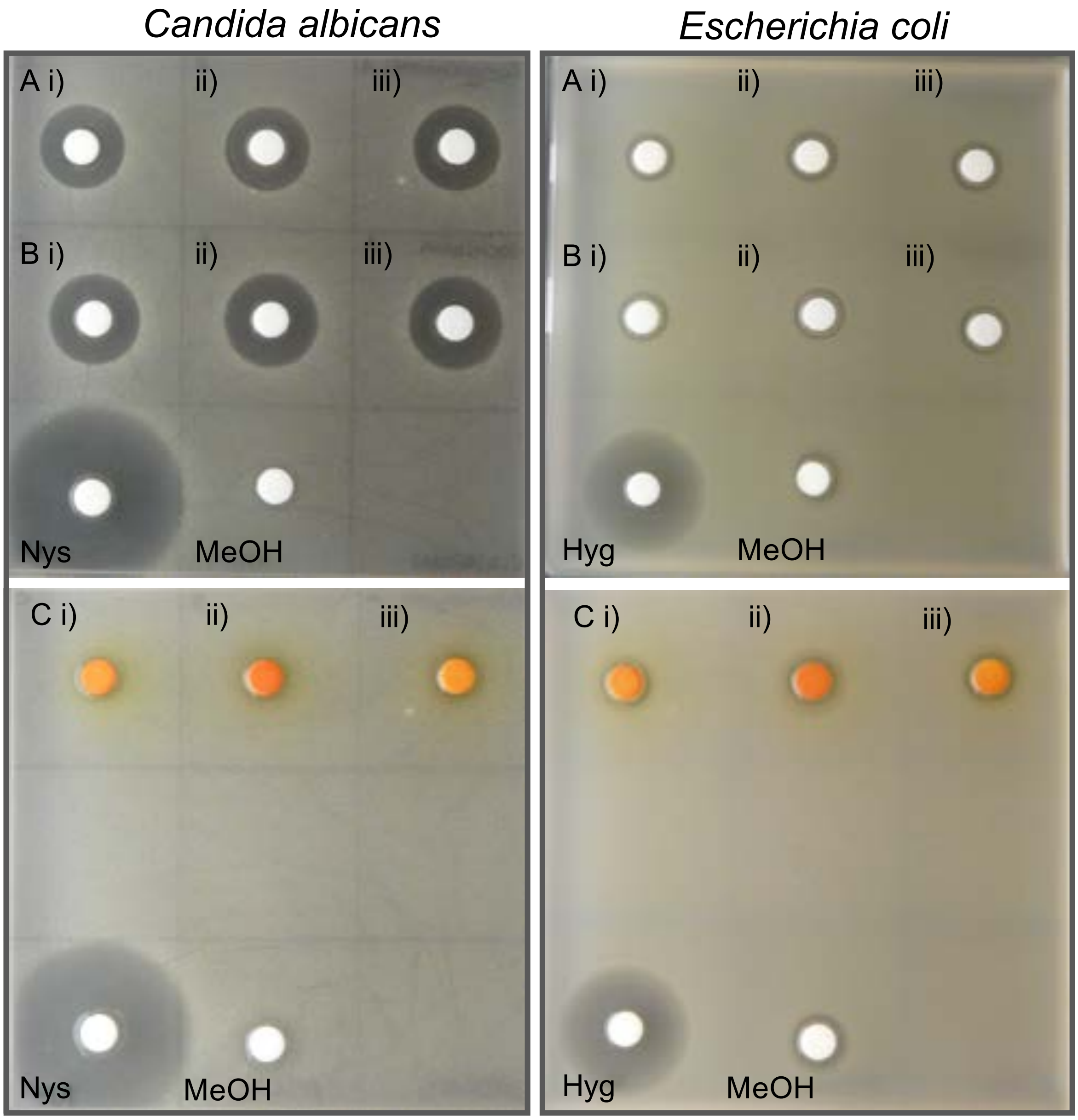

**Figure S18.** Disc-diffusion bioassays using compounds purified from the *Streptomyces* strain N2 against *Candida albicans* and *Escherichia coli*. Discs were soaked in purified extracts of either A) pentamycin, B) 14-hydroxyisochainin or C) actinomycin complex. N=3 (i-iii) technical replicates of each extract. Nystatin (Nys) or hygromycin (Hyg) were used as positive controls and methanol (MeOH) was used as a negative control.

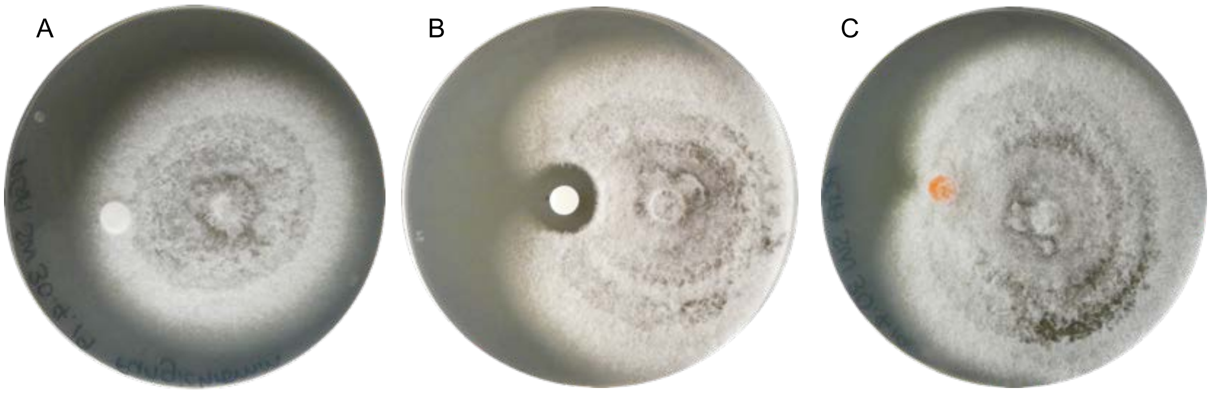

**Figure S19.** Disc-diffusion bioassays using compounds purified from *Streptomyces* strain N2 against *G. graminis* var. *tritici* (Take-all fungus). Discs were soaked in purified extracts of either: A) pentamycin: B) 14-hydroxyisochainin; or C) actinomycin complex. Antifungal activity is indicated by a zone of clearing around the disc.

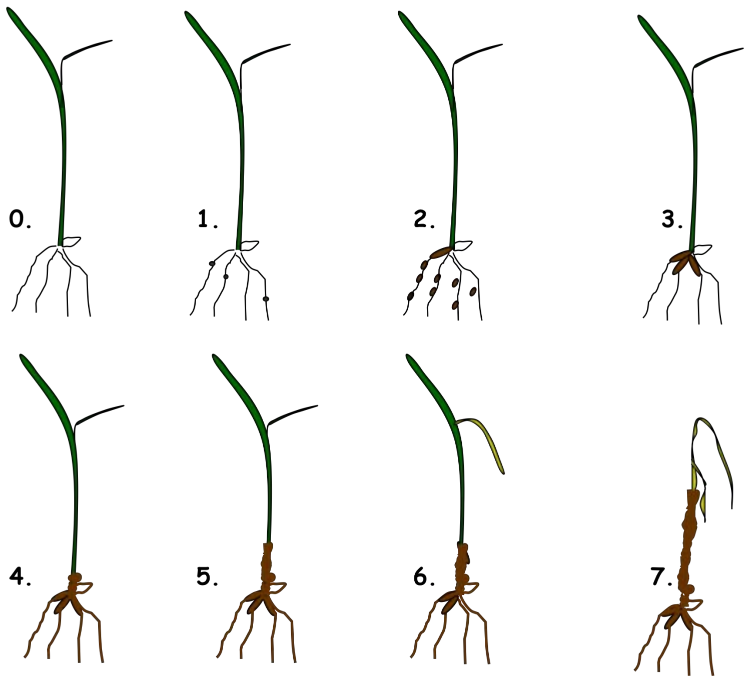

***Gaeumannomyces graminis* infection scoring system**

0. No infection
1. Very few lesions – only one per root
2. More than one lesion per root
3. Many small lesions and a few large lesions
4. Many large lesions
5. Roots almost completely brown
6. Roots mainly brown and evidence of infection in stem.
7. Roots brown, infected stem, leaves becoming yellow and wilted.
8. Entire plant brown and wilted.

**Figure S20.** Infection scoring system for the vermiculite-based Take-all infection assay of *Triticum aestivum.*

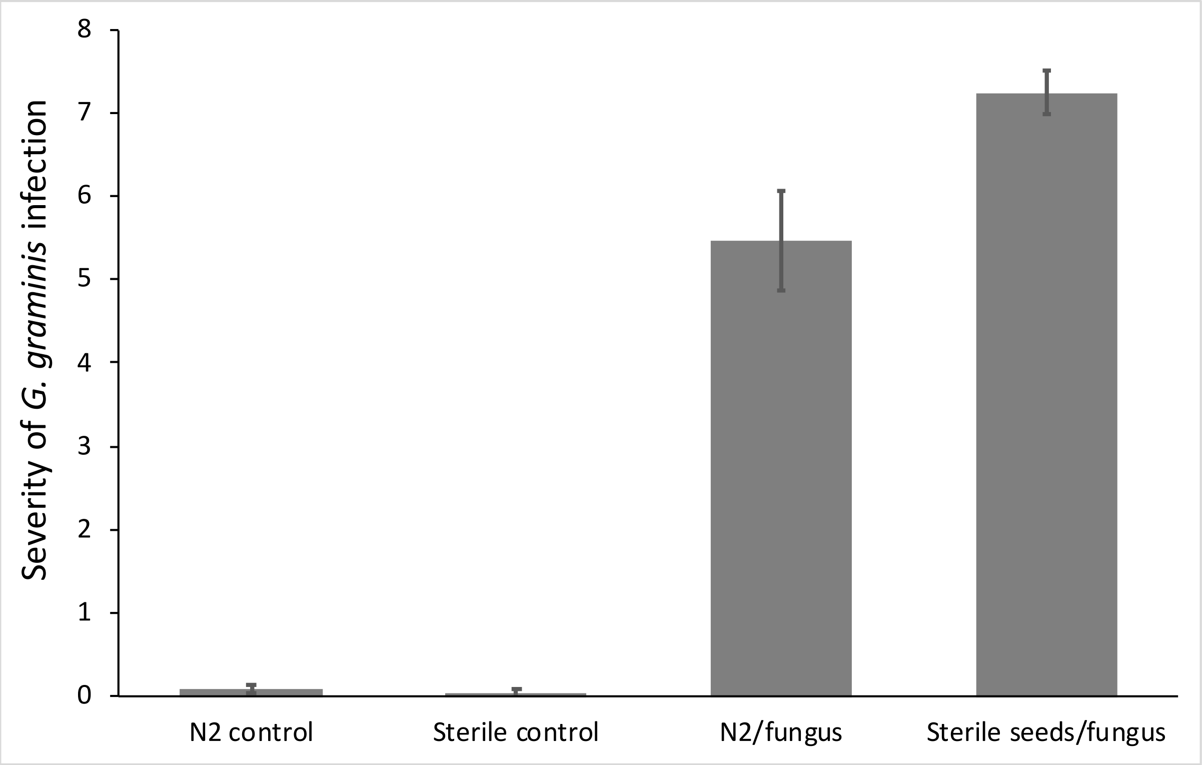

**Figure S21.** The effect of *Streptomyces* strain N2 on wheat plant infection severity by *G. graminis* var. *tritici*. Infections were scored after three weeks of growth. N2 control= seeds coated in N2 spores/no *G. graminis*; Sterile control= sterile seeds/no *G. graminis*; N2/fungus= seeds coated in N2 spores grown in the presence of *G. graminis*; Sterile seeds/fungus= sterile seeds grown in the presence of *G. graminis*. N=25 plants per treatment group, error bars represent standard errors.

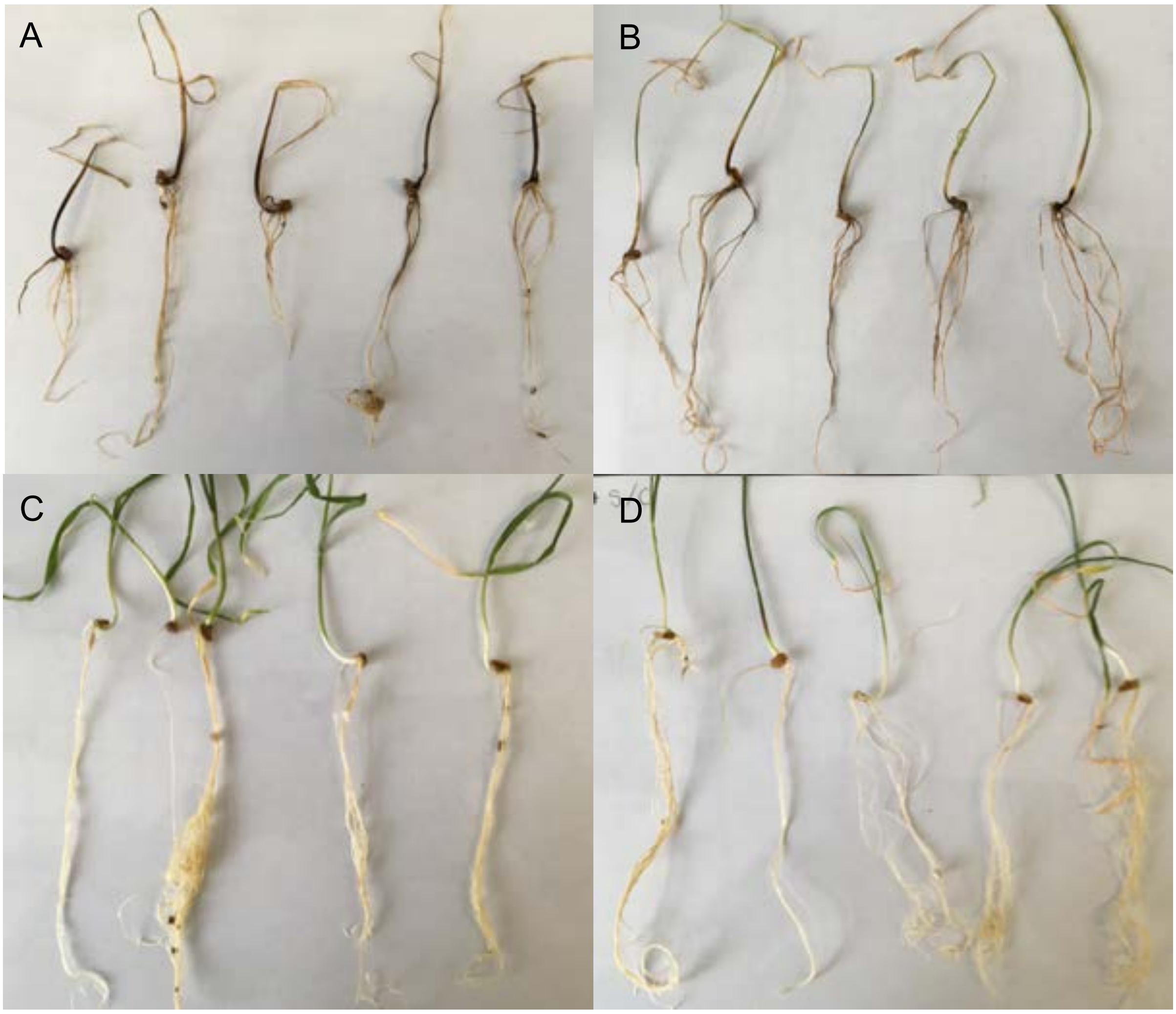

**Figure S22.** Wheat plants were grown A) from sterile seeds in the presence of *G. graminis* var. *tritici*, B) from seeds inoculated with N2 spores, in the presence of *G. graminis* var. *tritici* C) from sterile seeds, no *G. graminis* var. *tritici* D) from seeds coated with N2 spores, no *G. graminis* var. *tritici*.

**Supplementary tables**

**Table S1.** Strains, primers and plasmids used in experiments.

| Species/strain name | Description | Origin | Genome accession number |
|---|---|---|---|
| *Streptomyces* L2 | Wild-type | *A.thaliana* root microbiome, this study | QBDT00000000 |
| *Streptomyces* M2 | Wild-type | *A.thaliana* root microbiome, this study | CP028834 |
| *Streptomyces* M3 | Wild-type | *A.thaliana* root microbiome, this study | QANR00000000 |
| *Streptomyces* N1 | Wild-type | *A.thaliana* root microbiome, this study | QBDS00000000 |
| *Streptomyces* N2 | Wild-type | *A.thaliana* root microbiome, this study | CP028719 |
| *Streptomyces lydicus* ATCC25470 | Wild-type | American Type Culture Collection | RDTD00000000 |
| *Streptomyces lydicus* ATCC31975 | Wild-type | American Type Culture Collection | RDTE00000000 |

| | | | |
|---|---|---|---|
| *Streptomyces lydicus* Actinovate | Wild-type | Isolated from Actinovate™ by Elaine Patrick, UEA | RDTC00000000 |
| *Bacillus subtilis* | Wild-type, strain 168 | Gift from Nicola Stanley-Wall, University of Dundee | NA |
| Methicillin resistant *Staphylococcus aureus* | Clinical isolate | Norfolk and Norwich University Hospital (UK) | NA |
| *Escherichia coli* K12 | Wild-type | Lab stock, UEA | NA |
| *Pseudomonas syringae* DC3000 | Wild-type | John Innes Centre, Norwich, UK | NA |
| *Candida albicans* | Clinical isolate | Gift from Neil Gow, University of Exeter | NA |
| *Lomentospora prolificans* | Environmental isolate | American Type Culture Collection | NA |
| *Gaeumannomyces graminis* var. *tritici* | Environmental isolate | John Innes Centre, Norwich, UK | NA |
| *Arabidopsis thaliana* Col-0 | Wild-type, ecotype Col-0 | Lab stock, UEA | NA |
| *Triticum aestivum* var. Paragon | Wild-type, var. Paragon | John Innes Centre, Norwich, UK | NA |

| **Primer name** | **Sequence** | **Reference** |
|---|---|---|
| PRK341F | 5’-CCTACGGGAGGCAGCAG-3’ | Yu et al 2005 |
| MPRK806R | 5’-GGACTACHVGGGTWTCTAAT-3’ | |
| **Plasmid name** | **Description** | **Reference** |
| pIJ8660 | ermEp* driving constitutive production of codon optimised eGFP | Sun et al 1999 |

**Table S2.** The presence and absence of genes encoding enzymes involved in plant growth promotion pathways in sequenced *Streptomyces* isolates. Green indicates the presence of the enzyme in a given isolate, red indicates the absence of the enzyme.

| **Pathway** | **Enzyme** | **KO number** | **Kegg Reaction** | ***Streptomyces* strain** | | | | | | | | |
|---|---|---|---|---|---|---|---|---|---|---|---|---|
| | | | | **MG** | **M2** | **M3** | **N1** | **N2** | **L2** | ***S. lydicus* 25470** | ***S. lydicus* 31975** | **Actinovate** |
| IAA biosynthesis: IpyA pathway | Tryptophan aminotransferase | K14265 | R00684 | | | | | | | | | |
| | Indole-3-pyruvate decarboxylase | K04103 | R01974 | | | | | | | | | |
| | Acetaldehyde oxidase/dehydrogenase | K11817/ K00128 | R02681/ R02678 | | | | | | | | | |
| IAA biosynthesis: IAM pathway | Tryptophan 2-monooxygenase (IaaM) | K00466 | R00679 | | | | | | | | | |
| | IAM hydrolase (IaaH) | K01426/ K21801 | R03096 | | | | | | | | | |
| IAA biosynthesis: IAN pathway | Nitrilase | K01501 | R03093 | | | | | | | | | |
| IAA biosynthesis: TAM pathway | Tryptophan decarboxylase | K01593 | R00685 | | | | | | | | | |
| | (Mono or di) Amine oxidase | K00274/ K11182 | R02173 | | | | | | | | | |
| | Acetaldehyde oxidase/dehydrogenase | K11817/ K00128 | R02681/ R02678 | | | | | | | | | |

| | | | |
|---|---|---|---|
| Regulation of plant ethylene production | 1-aminocyclopropane-1-carboxylic acid (ACC) deaminase | K01505 | R00997 |

**Table S4.** Bioactivity screens of plant-associated *Streptomyces* species (purple columns, top) against a range of different indicator species (blue column, down). Screens were conducted on a range of different media (see Table S5 for recipes). Green squares indicate inhibition of the indicator strain by the given streptomycete, red indicates no inhibition.

| **Pathogen strain** | **Media** | **Streptomycete strain** | | | | | | | |
|---|---|---|---|---|---|---|---|---|---|
| | | **N1** | **N2** | **M3** | **M2** | **L2** | ***S. lydicus* 25470** | ***S. lydicus* 31975** | **Actinovate** |
| ***Bacillus subtilis*** | SFM | | | | | | | | |
| | Minimal salts | | | | | | | | |
| | ISP2 | | | | | | | | |
| | Oatmeal | | | | | | | | |
| | MYM | | | | | | | | |
| ***Methicillin-resistant Staphylococcus aureus*** | SFM | | | | | | | | |
| | Minimal salts | | | | | | | | |
| | ISP2 | | | | | | | | |
| | Oatmeal | | | | | | | | |
| | MYM | | | | | | | | |
| ***Escherichia coli*** | SFM | | | | | | | | |
| | minimal salts | | | | | | | | |
| | ISP2 | | | | | | | | |
| | Oatmeal | | | | | | | | |

| | |
|---|---|
| | MYM |
| ***Pseudomonas syringae DC3000*** | SFM |
| | Minimal salts |
| | ISP2 |
| | Oatmeal |
| | MYM |
| ***Candida albicans*** | SFM |
| | Minimal salts |
| | ISP2 |
| | PGA |
| | Oatmeal |
| | MYM |
| ***Gaeumannomyces graminis*** | PGA |
| ***Lomentospora prolificans*** | PGA |

**Table S5.** NMR shift data for pentamycin; 600/150 MHz in DMSO-$d_6$. *Signals may interchange due to resonance.

| Position | $^{13}C$ | $^{1}H$ (HSQC) | HMBC | COSY |
|---|---|---|---|---|
| **1** | 171.08 | | 2;3;27 | |
| **16** | 138.84 | | 15;8;29 | |
| **25** | 135.11 | 6.04 (dd, J=14.6 Hz; 4.5 Hz) | 23;24;26;27 | 24;26 |
| **19-23*** | 133.32 | 6.40 (m) | | |
| **19-23*** | 133.26 | 6.34 (m) | | |
| **19-23*** | 133.07 | 6.25 (m) | | |
| **19-23*** | 132.32 | 6.26 (m) | | |
| **19-23*** | 131.30 | 6.25 (m) | | |
| **24** | 129.08 | 6.34 (m) | 26 | 25 |
| **18** | 127.94 | 6.46 (dd, J=14.5 Hz, 11.4 Hz) | 17;29 | 17 |
| **17** | 127.14 | 5.94 (d, J=11.2 Hz) | 15;18;29 | 18 |
| **15** | 78.15 | 3.69 (d, J=8.9 Hz) | 14;17;29 | 14 |
| **14** | 76.56 | 3.48 (dd, J=8.9 Hz, 1.3 Hz) | 15 | 13;15 |
| **27** | 73.19 | 4.62 aqd (J=7.3 Hz, 6.4 Hz) | 25;26;28 | 26;28 |
| **26** | 71.16 | 3.98 (m) | 24;25;27;28 | 25;27 |
| **9** | 71.14 | 3.86 (m) | 8; 10 | 8;10 |
| **3** | 70.40 | 4.00 (m) | 2;4 | 2;4 |
| **7** | 70.29 | 3.89 (m) | 6; 8 | 6;8 |
| **5** | 70.00 | 3.89 (m) | 4 | 4;6 |
| **1´** | 69.58 | 3.67 (m) | 2 | 2;2´ |
| **11** | 69.30 | 3.77 (m) | 9;10;12;13 | 10;12 |
| **13** | 68.38 | 3.12 (d, J=10.7 Hz) | 14 | 12;14 |
| **2** | 58.73 | 2.46 (dd, J=8.5 Hz, 7.4 Hz) | 1´;2´;3;4 | 1´;3 |
| **8** | 44.02 ($CH_2$) | 1.39 (m), 1.28 (m) | 6;10 | 7;9 |
| **6** | 43.91 ($CH_2$) | 1.34 (m), 1.30 (m) | 4;8 | 5;7 |
| **10** | 42.93 ($CH_2$) | 1.36 (m), 1.26 (m) | 8;12 | 9;11 |
| **4** | 40.15 ($CH_2$) | 1.38 (m) | 2;6 | 3;5 |
| **12** | 38.71 ($CH_2$) | 1.58 (m), 1.36 (m) | 10;11;13;14 | 11;13 |
| **2´** | 34.19 ($CH_2$) | 1.35 (m), 1.25 (m) | 1´;2;3´;4´ | 1´ |
| **4´** | 31.25 ($CH_2$) | 1.25 (m), 1.19 (m) | 2´;3´;5´;6´ | |
| **3´** | 24.59 ($CH_2$) | 1.43 (m), 1.25 (m) | 1´;2´;4´ | |
| **5´** | 22.12 ($CH_2$) | 1.26 (m) | 4´;6´ | 6´ |
| **28** | 17.80 | 1.19 (d, J=6.3 Hz) | 26;27 | 27 |
| **6´** | 13.96 | 0.85 (t, J=7.0 Hz) | 5´ | 5´ |
| **29** | 11.67 | 1.69 (s) | 15;17 | 17 |

**Table S6.** NMR data for 14-hydroxyisochainin; 600/150 MHz in DMSO-$d_6$. *Signals can interchange due to resonance; **tentative.

| Position | $^{13}C$ | $^{1}H$ (HSQC) | HMBC | COSY |
|---|---|---|---|---|
| **1** | 173.19 | | 1´;2;3;27 | |
| **16** | 138.92 | | 15;18;29 | |
| **25** | 135.08 | 5.96 (m) | 24;26;27 | 24;26 |
| **19-23*** | 133.04 | 6.32 (m) | | |
| **19-23*** | 132.98 | 6.32 (m) | | |
| **19-23*** | 132.85 | 6.32 (m) | | |
| **19-23*** | 132.45 | 6.26 (m) | | |
| **19-23*** | 131.63 | 6.25 (m) | | |
| **24** | 129.12 | 6.33 (m) | 26 | 23;25 |
| **18** | 128.01 | 6.46 (dd, J=11.7, 11.1 Hz); | 17;29 | 17 |
| **17** | 127.05 | 5.94 (m) | 15;18;29 | 18;29 |
| **15** | 78.02 | 3.69 (d, J=8.8 Hz) | 14;17;29 | 14 |
| **14** | 76.53 | 3.48 (dd, J=8.9; 1.2 Hz) | 15 | 13;15 |
| **27** | 72.99 | 4.63 (m) | 26;28 | 26;28 |
| **26** | 71.31 | 3.95 (dd, J= 8.4 Hz; 4.7 Hz) | 24;25;27;28 | 25;27 |
| **9** | 71.12 | 3.86 (m) | 8;10 | |
| **3** | 70.85 | 3.61 (m) | 2;4 | 2;4 |
| **7**** | 69.64 | 3.88 (m) | 6;8 | 6 |
| **5**** | 69.51 | 3.88 (m) | 4;6 | 6 |
| **11** | 69.28 | 3.79 (m) | 10;12;13 | 10;12 |
| **13** | 68.23 | 3.13 (d, J=10.8 Hz) | 12;14 | 12;14 |
| **2** | 52.49 | 2.25 (m) | 1´;2´;3;4 | 1´;3 |
| **8** | 43.84 ($CH_2$) | 1.39 (m) | 6;10 | 7 |
| **6** | 43.56 ($CH_2$) | 1.31 (m) | 4;8 | 5;7 |
| **10** | 42.60 ($CH_2$) | 1.37 (m), 1.27 (m) | 8;12 | 11 |
| **4** | 41.61 ($CH_2$) | 1.31 (m) | 2;6 | 3 |
| **12** | 38.68 ($CH_2$) | 1.58 (m), 1.36 (m) | 10;14 | 11;13 |
| **1´** | 28.82 ($CH_2$) | 1.67 (m), 1.43 (m) | 2;2´;3;3´; | 2;2´ |
| **2´** | 28.82 ($CH_2$) | 1.17 (m) | 1´;2;3´;4´ | 1´ |
| **3´** | 22.00 ($CH_2$) | 1.26 (m) | 2´;4´ | 4´ |
| **28** | 18.07 | 1.22 (d, J=6.4 Hz) | 26 | 27 |
| **4´** | 13.77 | 0.84 (t, J=7.2 Hz) | 2´; 3´ | 3´ |
| **29** | 11.76 | 1.69 (s) | 17 | |

**Table S7.** Comparison of $^{13}C$ chemical shifts measured at 600 MHz in $CD_3OD$ for 14-hydroxyisochainin isolated in this study and reported in Li *et al* (52).

| $^{13}C$ shifts this study | Position | $^{13}C$ shifts Li *et al* |
|---|---|---|
| 175.40 | 1 | 175.37 |
| 138.68 | 16 | 138.71 |
| 135.17 | 19 | 135.18 |
| 134.44 | 21 | 134.45 |
| 134.36 | 25 | 134.37 |
| 134.26 | 23 | 134.32 |
| 133.95 | 20 | 133.96 |
| 133.72 | 22 | 133.74 |
| 132.01 | 24 | 131.99 |
| 129.80 | 17 | 129.79 |
| 129.24 | 18 | 129.25 |
| 80.32 | 15 | 80.32 |
| 78.20 | 14 | 78.20 |
| 74.58 | 27 | 74.58 |
| 73.97 | 9 | 74.02 |
| 73.59 | 5 | 73.64 |
| 73.58 | 7 | 73.56 |
| 73.48 | 3 | 73.55 |
| 73.37 | 26 | 73.44 |
| 71.34 | 11 | 71.35 |
| 70.27 | 13 | 70.26 |
| 54.40 | 2 | 54.40 |
| 45.15 | 8 | 45.16 |
| 44.82 | 6 | 44.83 |
| 44.13 | 10 | 44.15 |
| 42.34 | 4 | 42.33 |
| 39.49 | 12 | 39.50 |
| 30.56 | 1´ | 30.57 |
| 30.16 | 2´ | 30.18 |
| 23.59 | 3´ | 23.61 |
| 18.30 | 28 | 18.30 |
| 14.24 | 4´ | 14.25 |
| 11.80 | 29 | 11.80 |

**Table S8.** Media recipes used in experiments

| Media | Component | g $L^{-1}$ $dH_2O$ |
|---|---|---|
| Soya Flour Mannitol (SFM) agar (Kieser et al, 2000) | Soy flour | 20 |
| | Mannitol | 20 |
| | Agar | 20 |
| Minimal Salts medium (Lebeis et al, 2012) | $NH_4SO_4$ | 2 |
| | $K_2HPO_4$ | 14 |
| | $KH_2PO_4$ | 6 |
| | Sodium citrate | 1 |
| | $MgSO_4$ | 0.2 |
| | Agar | 15 |
| Maltose-Yeast extract-Malt extract (MYM) agar | Maltose | 4 |
| | Yeast Extract | 4 |
| | Malt Extract | 10 |
| | Agar | 18 |
| ISP2 Agar | Yeast Extract | 4 |
| | Maltose | 10 |
| | D-glucose | 4 |
| | Agar | 20 |
| Oatmeal Agar | Ground oats | 20 |
| | Agar | 20 |
| Potato Glucose Agar (PGA) | PGA (Sigma Aldrich) | 39 |
| Lysogeny Broth (LB) | Tryptone | 10 |
| | NaCl | 10 |
| | Yeast extract | 5 |

| | | |
|---|---|---|
| | Glucose (only for growth of *P. syringae*) | 1 |
| Minimal medium ± IAA (Kieser et al, 2000) | $NH_4SO_4$ | 1 |
| | $KH_2PO_4$ | 0.5 |
| | $MgSO_4.7H_2O$ | 0.2 |
| | $FeSO_4.H_2O$ | 0.01 |
| | Agar | 15 |
| | Trace elements (added after autoclaving) | 2 ml |
| | ± Indole Acetic Acid (IAA, Sigma Aldrich) | 0.1 |
| Dworkins and Foster medium (Dworkin and Foster, 1958) | $(NH_4)_2SO_4$ | 2 |
| | $KH_2PO_4$ | 4 |
| | $Na_2HPO_4$ | 6 |
| | $MgSO_4.7H_2O$ | 0.2 |
| | $FeSO_4.H_2O$ | 0.001 |
| | $H_3BO_4$ | 0.0001 |
| | $MnSO_4$ | 0.0001 |
| | $ZnSO_4$ | 0.0007 |
| | $CuSO_4$ | 0.0005 |
| | $MoO_3$ | 10 |
| | Agar | 20 |
| 2xYT | Bacto-tryptone | 16 |
| | Yeast extract | 10 |
| | NaCl | 5 |

| | | |
|---|---|---|
| Murashige and Skoog (MSk) Agar | Murashige and Skoog salts (Duchefa Biochemie, Harlem, Netherlands) | 4.43 |
| | Sucrose | 10 or 0 |
| | Agar | 8 or 15 |
| Yeast Peptone Dextrose (YPD) Agar | Yeast Extract | 10 |
| | Bactopeptone | 40 |
| | Glucose | 15 |
| | Agar | 15 |
| Silwett L-77 amended Phosphate Buffered Saline (PBS-S) | $NaH_2PO_4.H_2O$ | 6.33 g |
| | $Na_2HPO_4.H_2O$ | 16.5 g |
| | 200 µl Silwet L-77 added after autoclaving | |
